# Structural basis for potent antibody neutralization of SARS-CoV-2 variants including B.1.1.529

**DOI:** 10.1101/2021.12.27.474307

**Authors:** Tongqing Zhou, Lingshu Wang, John Misasi, Amarendra Pegu, Yi Zhang, Darcy R. Harris, Adam S. Olia, Chloe Adrienna Talana, Eun Sung Yang, Man Chen, Misook Choe, Wei Shi, I-Ting Teng, Adrian Creanga, Claudia Jenkins, Kwanyee Leung, Tracy Liu, Erik-Stephane D. Stancofski, Tyler Stephens, Baoshan Zhang, Yaroslav Tsybovsky, Barney S. Graham, John R. Mascola, Nancy J. Sullivan, Peter D. Kwong

## Abstract

With B.1.1.529 SARS-CoV-2 variant’s rapid spread and substantially increased resistance to neutralization by vaccinee and convalescent sera, monoclonal antibodies with potent neutralization are eagerly sought. To provide insight into effective neutralization, we determined cryo-EM structures and evaluated potent receptor-binding domain (RBD) antibodies for their ability to bind and neutralize this new variant. B.1.1.529 RBD mutations altered 16% of the RBD surface, clustering on a ridge of this domain proximal to the ACE2-binding surface and reducing binding of most antibodies. Significant inhibitory activity was retained, however, by select monoclonal antibodies including A19-58.1, B1-182.1, COV2-2196, S2E12, A19-46.1, S309 and LY-CoV1404, which accommodated these changes and neutralized B.1.1.529 with IC_50_s between 5.1-281 ng/ml, and we identified combinations of antibodies with potent synergistic neutralization. Structure-function analyses delineated the impact of resistance mutations and revealed structural mechanisms for maintenance of potent neutralization against emerging variants.

**Summary Sentence:** We show potent B.1.1.529 neutralization by select antibodies and use EM structures to reveal how potency can be retained.

## Introduction

Since first appearing in late 2019 (*1*), SARS-CoV-2 has infected over 280 million people and resulted in over 5.4 million deaths (*2*). In the summer of 2020 when the B.1.1.7 (Alpha) variant was first reported, it became apparent that host-virus interactions and virus adaption to immune pressures were leading to the generation of variants with variable pathogenicity, transmissibility and increasingly, the ability to escape from vaccines and therapeutic antibody treatments. The appearance and rapid spread of the B.1.1.529 (Omicron) variant (*3, 4*), with almost 40 mutations in spike, three times higher than the number found in prior variants, has raised alarm. How do these mutations impact spike? How do they alter recognition by antibody? And can potent neutralization be maintained? While extremely broad antibodies such as S2P6 (*5*) have been identified that neutralize diverse beta-coronaviruses including SARS-CoV-2 and are likely to be unencumbered by B.1.1.529 mutations, these broad antibodies neutralize in the microgram per ml range, whereas current therapeutic antibodies generally neutralize ~100-times more potently, in the 1-50 nanogram per ml range for the ancestral D614G virus. In this study, we determine cryo-EM structures of the B.1.1.529 spike, measure the binding and neutralization capacity of leading monoclonal antibodies against five variants of concern (VOC), including B.1.1.529, and use functional and structural analyses to determine the bases for differences in variant neutralization and to identify antibodies and combinations of antibodies capable of potently neutralizing emerging variants including B.1.1.529.

## Cryo-EM structure of B.1.1.529 (Omicron) spike

To provide insight into the impact of B.1.1.529 mutations on spike, we expressed and produced the two proline-stabilized (S2P) (*6*) B.1.1.529 spike and collected single particle cryo-EM data to obtain a structure of the trimeric ectodomain at 3.29 Å resolution (**Fig. 1A, Fig. S1, Fig S2 and Table S1**). Like other D614G containing variants, the most prevalent spike conformation comprised the single-receptor-binding domain (RBD)-up conformation (*7*). B.1.1.529 mutations present in the spike gene resulted in 3 deletions of 2, 3 and 1 amino acids, a single insertion of 3 amino acids and 30 amino acid substitutions in the spike ectodomain. As expected from the ~3% variation in sequence, the B.1.1.529 spike structure was extremely similar to the WA-1 spike structure with an overall Cα-backbone RMSD of 1.8 Å (0.5 Å for the S2 region); however, we did observe minor conformational changes in a few places. For example, the RBD S371L/S373P/S375F substitutions changed the conformation of their residing loop, with Phe375 in the RBD-up protomer formed Phe-Phe interaction with Phe486 in the neighboring RBD-down protomer (**Fig. 1B**), helping to stabilize the single-RBD-up conformation. Amino acid changes were denser in the N-terminal domain (NTD) and RBD, where a majority of neutralization occurs, though RMSDs remained low (0.6 Å and 1.2 Å for NTD and RBD, respectively). Notably, about half the B.1.1.529 alterations in sequence outside NTD and RBD involved new interactions, both hydrophobic, such as Tyr796 with glycan on Asn709, and electrostatic, such as Lys547 and Lys856 interacting with residues in HR1 SD1 on neighboring protomers (**Fig. 1B, Table S2**). These heightened interprotomer interactions suggested a need to maintain trimer stability. Differential scanning calorimetry indicated the B.1.1.529 spike to have folding energy similar to the original WA-1 strain (**Fig. S2**).

**Figure 1.**
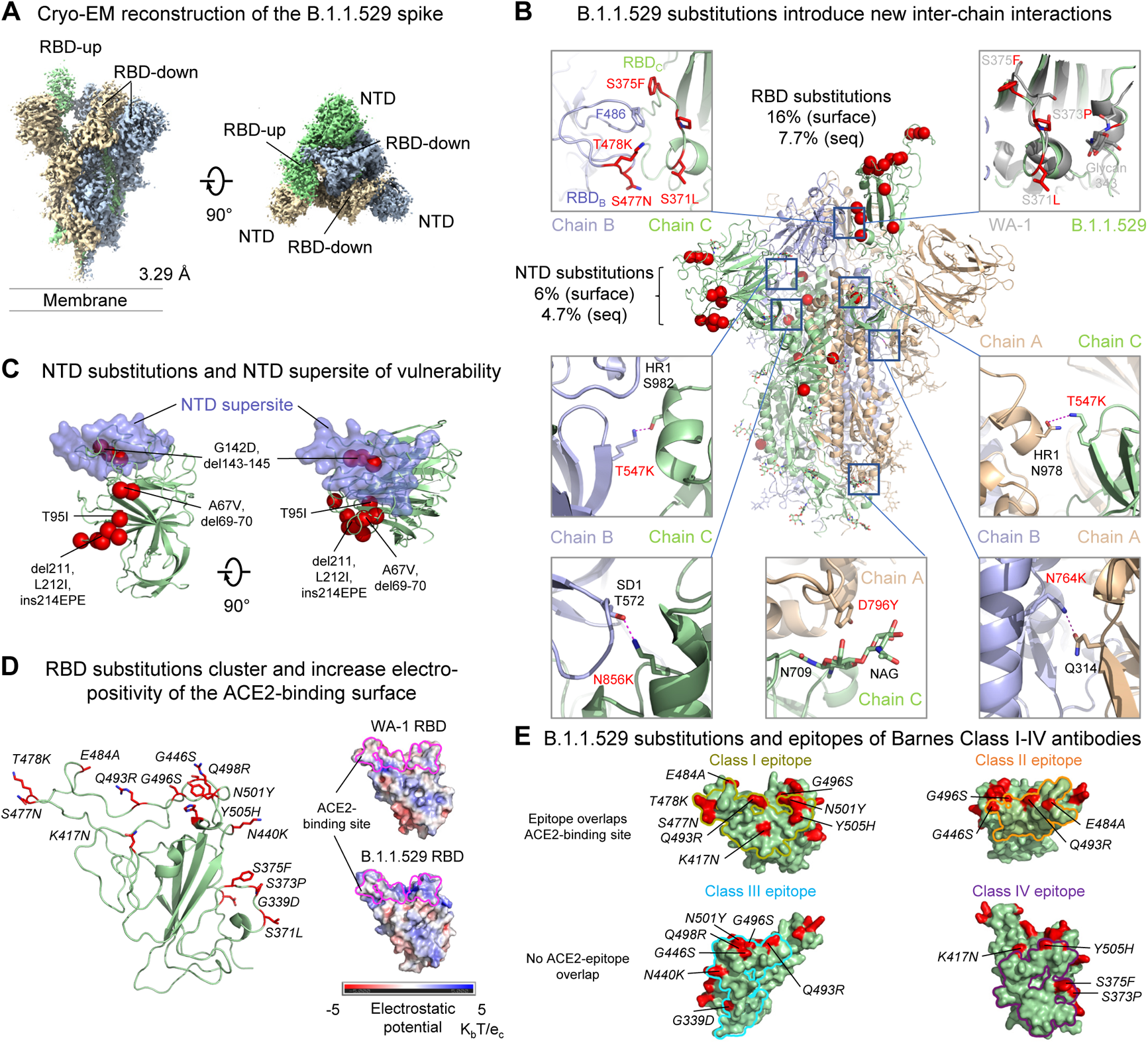
Cryo-EM structure of the SARS-CoV-2 B.1.1.529 (Omicron) spike. A. Cryo-EM map of the SARS-CoV-2 B.1.1.529 spike. Reconstruction density map at 3.29 Å resolution is shown with side and top views. Protomers are colored light green, wheat and light blue. The contour level of cryo-EM map is 4.0s.
B. B.1.1.529 amino acid substitutions introduced inter-protomer interactions. Substitutions in one protomer are shown as red spheres. Examples of inter-protomer interactions introduced by B.1.1.529 substitutions were highlighted in box with zoom-in view to the side. Mutation are described as a percentage of the domain surface (surface) or as a percentage of the sequence (seq).
C. The NTD supersite of vulnerability is shown in semi-transparent surface along with a green backbone ribbon. Amino acid substitutions, deletions, and insertions are colored red.
D. The 15 amino acid substitutions clustered on the rim of RBD, changed 16 % of the RBD surface areas (left) and increased electro-positivity of the ACE2-binding site (right). Mutated residues were shown in red sticks. The ACE2-binding site on the electrostatic potential surface were marked as magenta lines.
E. Mapping B.1.1.529 RBD substitutions on the epitopes of Barnes Class I-IV antibodies. The locations of the substitutions were shown in red on the surface. Those may potentially affect the activity of antibodies in each class were labeled with their residue numbers. Class I footprint were defined by epitopes of CB6 and B1-182.1, Class II footprint were defined by epitopes of A19-46.1 and LY-CoV555, Class III footprint were defined by epitopes of A19-61.1, COV2-2130, LY-CoV1404 and S309, Class IV footprint were defined by epitopes of DH1407 and S304. Class I and II epitope have overlap with the ACE2 binding site, while class III and IV do not. Class II and III epitopes allow binding to WA-1 when RBD is in the up or down conformation.

NTD changes altered ~6% of the solvent accessible surface on this domain, and several were located directly on or proximal to the NTD-supersite of vulnerability (*8*), where prior variants had mutations that substantially reduced neutralization by NTD antibodies. Other NTD changes were proximal to a pocket, proposed to be the site of bilirubin binding (*9*), which also binds antibody (*10*) (**Fig. 1C**).

RBD alterations changed ~16% of the solvent accessible surface on this domain and were constrained to the outward facing ridge of the domain (**Fig. 1D**), covering much of the surface of the trimeric spike apex (**Fig. S1F**). Several amino acid changes involved basic substitutions, resulting in a substantial increase in RBD electro-positivity (**Fig. 1D**). Overall, RBD changes were located proximal to binding surfaces for the ACE2 receptor (*11*) (**Fig. 1D**) as well as to recognition sites for potently neutralizing antibodies (**Fig. 1E**) (*12–14*).

## Functional assessment of variant binding to ACE2

When pathogens infect a new species, sustained transmission leads to adaptations that optimize replication, immune-avoidance and transmission. One hypothesis for the efficient species adaptation and transmission of SARS-CoV-2 in humans is that the virus spikes are evolving to optimize binding to the host receptor protein, ACE2. As a first test of this hypothesis, we used a flow cytometric assay to evaluate binding of human ACE2 to cells expressing variant spike proteins. We evaluated the binding of soluble dimeric ACE2 to B.1.1.7 (*15*), B.1.351 (Beta) (*16*), P.1 (Gamma) (*17, 18*) or B.1.617.2 (Delta) (*19*) spikes compared to the ancestral D614G spike. The earliest variants, B.1.1.7, B.1.351 and P.1, first recognized in December 2020 and January 2021 contain an RBD mutation at N501Y, which has been proposed to increase binding to ACE2 (*20*). Cell surface ACE2 binding to B.1.1.7 was 124% of D614G, while B.1.351, and P.1 were 69% and 76%, respectively (**Fig S3A**), demonstrating that these variants do not have a substantial increase in ACE2 affinity. We next evaluated binding to the B.1.1.529 spike protein and observed ACE2 binding to be 200% that of D614G variant (**Fig S3A**).

Since cell-surface binding may involve other factors, such as might be impacted by the increased electro-positivity of the RBD noted above, and formal kinetic measurements are challenging to obtain using cell-based assays, we further investigated ACE2 binding affinity using surface plasmon resonance measurements of soluble dimeric human ACE2 to S2P spike trimers generated from the ancestral WA-1 and 6 subsequent variants: D614G, B.1.351, P.1, B.1.617.2, B.1.1.7 and B.1.1.529. We observed that both WA-1 and D614G, which have identical RBD sequences, have similar affinities (*K_D_*= 1.1 nM and 0.73 nM, respectively) (**Fig S3B,C**). We noted that the affinity for B.1.617.2, which contains two RBD mutations, was ~3-fold worse (KD=2.4 nM) than the D614G S2P trimer. We next evaluated B.1.1.7, B.1.351, P.1 and B.1.1.529, which each contain N501Y substitutions, and found affinities of 0.59 nM, 1.7 nM, 0.85 nM and 3.8 nM, respectively (**Fig S3B,C**). Collectively, the cell surface and S2P binding results show minimal improved affinity in some, but not all spikes, and suggests that, while SARS-CoV-2 maintains nanomolar affinity to ACE2, spike variant evolution appears to be driven primarily by immune pressure.

## Variant binding and neutralization by individual monoclonal antibodies

To define of the impact SARS CoV-2 variant amino acid changes on the binding and neutralization of monoclonal antibodies, we expressed and purified 17 highly potent antibodies targeting the spike RBD (*12, 13, 21–33*); including 13 antibodies currently under clinical investigation or approved for use under expanded use authorization (EUA) by the United States Food and Drug Administration. All antibodies bound and neutralized B.1.1.7 comparable to the ancestral D614G and consistent with the single 501Y substitution being outside each antibody’s binding epitope (**Fig 2, Fig S4A**). The addition of two more RBD substitutions, K417N and E484K (**Fig 2A**) in the B.1.351 and P1 variants eliminated binding by two Class I antibodies, CB6 and REGN10933 and two Class II antibodies, LY-CoV555 and C144 (**Fig 2B, Fig S4A**). Neutralization of B.1.351 and P1 by CB6, LY-CoV555 and C144 was completely abolished, while REGN10933 was eliminated for B.1.351 and reduced >250-fold for P.1. In addition, while binding of CT-P59 to B.1.351 and P.1 variants was minimally changed (69-79%), neutralization was decreased 26-43-fold (IC_50_ 65.8 and 39.6 ng/mL) (**Fig 2B,C**). The remaining antibodies showed minimal binding changes and a <3.6-fold difference in neutralization IC_50_ (**Fig 2B,C, Fig S4A**). An evaluation of the antibodies in our panel against B.1.617.2 revealed minimal changes in binding for all antibodies except A19-46.1 and LY-CoV555, which were 0% of D614G (**Fig 2B**). Neutralization assays using B.1.617.2 pseudovirus showed that the REGN10987 IC_50_ was 22.7-fold lower than D614G and neutralization values for A19-46.1 and LY-CoV555 were each >10,000 ng/mL (**Fig 2C**). These data are consistent with previous results that showed both A19-46.1 and LY-CoV555 were sensitive to the L452R mutations present in B.1.617.2 (*14*).

**Figure 2.**
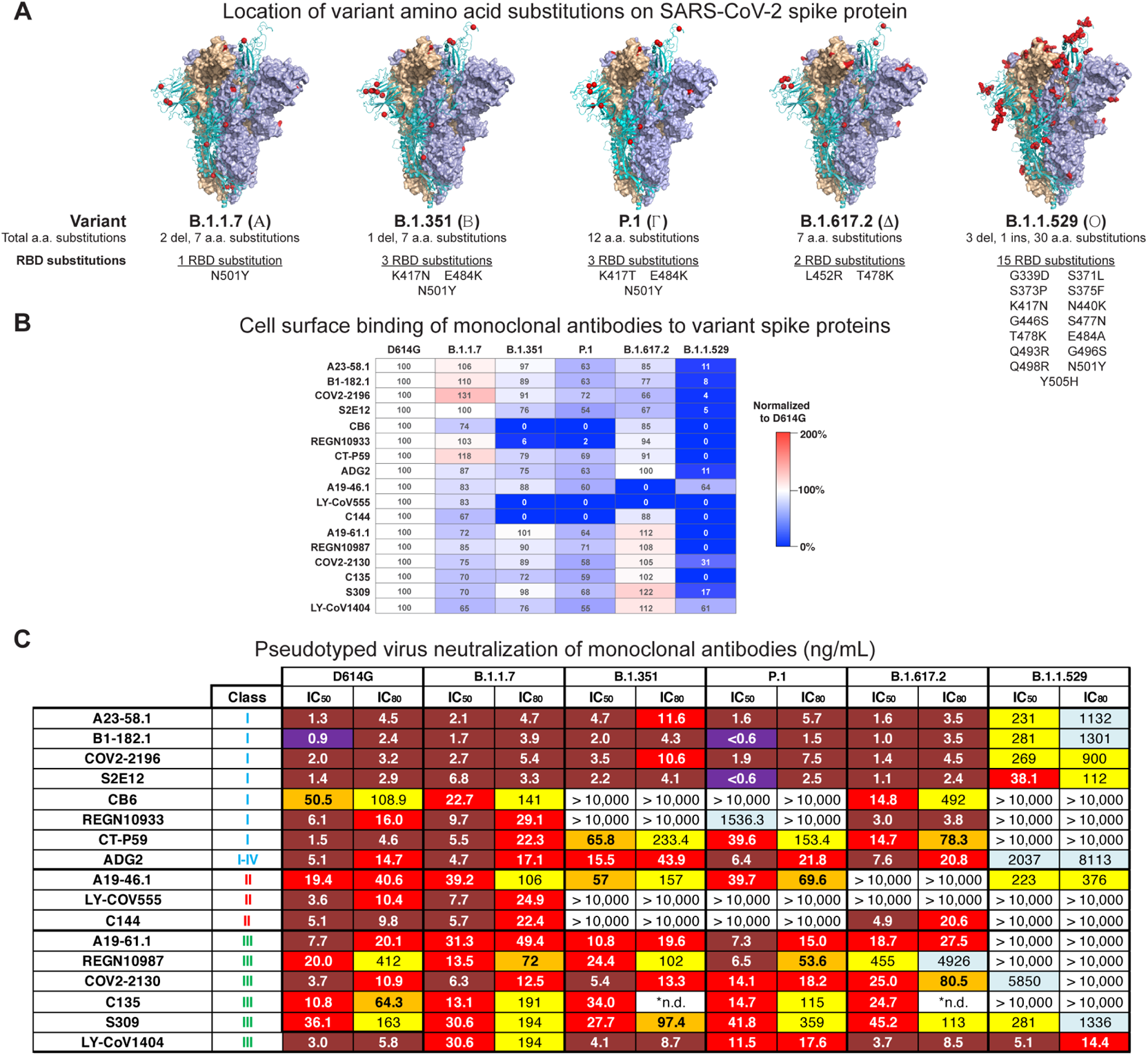
SARS-CoV-2 monoclonal antibody binding and neutralization. A. Models of SARS-CoV-2 WA-1 spike protein (PDB: 6XM3) with the locations of substitutions present of variants indicate as red dots. Also noted is the total number of mutation and the number and locations of receptor binding domain (RBD) mutants in variant of concern spike proteins.
B. Full length spike proteins from the indicated SARS-CoV-2 variants were expressed on the surface of transiently transfected 293T cells and binding to indicated monoclonal antibodies was assessed by flow cytometry. Shown is the mean fluorescence intensity (MFI) of bound antibody on the indicated cell relative to the MFI of the same antibody bound to D614G expressing cells. The data is expressed as a percentage. Shown is a representative experiment (n=2-3 for each antibody).
C. Lentiviruses pseudotyped with SARS-CoV-2 spike proteins from D614G, B.1.1.7, B.1.351, P.1, B.1.617.2 or B.1.1.529 were incubated with serial dilutions of the indicated antibodies and IC_50_ and IC_80_ values determined. S309 was tested on 293 flpin-TMPRSS2-ACE2 cells while all the other antibodies were tested on 293T-ACE2 cells. Ranges are indicated with white (>10,000 ng/ml), light blue (>1000 to ≤10,000 ng/ml), yellow (>100 to ≤1000 ng/ml), orange (>50 to ≤100 ng/ml), red (>10 to ≤50 ng/ml), maroon (>1 to ≤10 ng/ml), and purple (≤1 ng/ml). *n.d.= not determined due to incomplete neutralization that plateaued at <80% (See Fig S4B).

For B.1.1.529, we noted that all but three antibodies showed binding less than 31% of D614G. It is interesting to note that COV2-2196, S2E12, B1-182.1 and A23-58.1 utilize the same VH1-58 gene in their heavy chain and target a similar region on the RBD (i.e., VH1-58 supersite), but show differential binding to the B.1.1.529 (*i.e.,* 4%, 5%, 8% and 11%, respectively) and B.1.617.2 (*i.e.,* 66%, 67%, 77% and 85%, respectively) (**Fig 2B**). Even though the absolute differences in binding are minimal, the shared trend may be reflective of how the RBD tip mutation at T478K mutation is accommodated by each of these antibodies. Finally, LY-CoV1404 revealed 61% binding to B.1.1.529 spike. Taken together, cell surface binding suggests that while both A19-46.1 and LY-CoV1404 are likely to retain potent neutralizing activity against B.1.1.529, the remaining antibodies in our panel might show decreased neutralizing activity.

Using the same panel of monoclonal antibodies, we further assayed for each antibody’s capacity to neutralize the B.1.1.529 variant. While VH1-58 supersite antibodies (Class I) show high neutralization activity against other variants, antibodies targeting the supersite were 40 to 126-fold worse (IC_50_ 38-269 ng/ml) against B.1.1.529 viruses than D614G (IC_50_ 0.9-2.0 ng/ml) (**Fig 2C**). In addition, two other antibodies, CB6 (Class I) and ADG2 (Class I/IV) were shown to be severely impacted (IC_50_ >10,000 ng/mL CB6 and 2037 ng/mL ADG2 to B.1.1.529 vs 31 and 50.5 ng/mL to D614G, respectively) (**Fig 2C**). We next analyzed Class II antibodies (i.e., LY-CoV555, C144, A19-46.1) and found that amongst these, neutralization by LY-CoV555 and C144 was completely abolished (IC_50_ >10,000 ng/mL B.1.1.529 vs 3.6 and 5.1 ng/mL D614G, respectively). In contrast, we found that the A19-46.1 IC_50_ neutralization was 223 ng/mL for B.1.1.529 vs 19.4 ng/mL for D614G (**Fig 2C**) and was <6 fold of the previously reported IC_50_ for WA-1 (39.8 ng/mL) (*14*). We next analyzed the Class III antibodies (*i.e.,* A19-61.1, REGN10987, COV2-2130, C135, LY-CoV1404) and noted neutralization activity of A19-61.1, REGN10987 and C135 was completely abolished (IC_50_ >10,000 ng/mL B.1.1.529 vs 19.4, 20.0, 10.8 ng/mL, respectively on D614G), CoV2-2130 decreased 1581-fold (IC_50_ 5850 ng/mL B.1.1.529 vs 3.7 ng/mL D614G) and that of S309 decreased by ~8-fold (IC_50_ 281 ng/mL B.1.1.529 vs 36.1 ng/mL D614G) (**Fig 2C**). Strikingly, in contrast to all of the other antibodies, we found that the neutralization of LY-CoV1404 against B.1.1.529 was unchanged relative to D614G (IC_50_ 5.1 ng/mL for B.1.1.529 vs 3 ng/mL for D614G) (**Fig 2C**). Taken together, these data demonstrate that the mutations present in B.1.1.529 mediate resistance to antibodies.

## Structural and functional basis of Class I antibody neutralization, escape and retained potency

We sought to determine the functional basis of B.1.1.529 neutralization and escape for Class I antibodies and to understand how potent neutralization might be retained. We analyzed Class I antibodies, CB6, B1-182.1 and S2E12, with differential B.1.1.529 neutralization (**Fig 2C**). We first evaluated CB6 using virus particles containing single amino acid substitutions representing 13 of 15 single amino acid changes on the RBD of B.1.1.529 (*i.e*., all but S375F and G496S) or G496R; we noted while several minimally changed their neutralization IC_50_, only Y505H, S371L, Q493R and K417N decreased neutralization >5-fold, with IC_50_ of 50, 212, 320, and >10,000 ng/mL, respectively (**Fig 3A**). This suggests that B.1.1.529 evades CB6-like antibodies through multiple mutations. Docking of the RBD-bound CB6 onto the B.1.1.529 structure revealed several B.1.1.529 residues may potentially clash with CB6. Especially, K417N, Q493R and Y505H were positioned to cause severe steric hindrance to the CB6 paratope, consistent with the neutralization data (**Fig. 3B**). We next evaluated two VH1-58 supersite antibodies, B1-182.1 and S2E12, which have highly similar amino acid sequences but show ~6-fold difference in B.1.1.529 neutralizing. These two antibodies remained highly potent (<10.6 ng/mL IC_50_) for all virus particles with single RBD mutations, with the largest change for Q493R, which caused a 7 and 5.4-fold decrease of neutralization for B1-182.1 and S2E12, respectively. These small differences in neutralization from single mutations suggest that multiple mutations of B.1.1.529 are working in concert to mediate escape from VH1-58 supersite antibodies. Docking of the RBD-bound B1-182.1 onto the B.1.1.529 structure indicated that the epitopes of these VH1-58-derived antibodies were confined by Q493R, S477N, T478K and E484A (**Fig. 3C**). With R493 pressing on one side of the antibody like a thumb, N477/K478 squeezed onto the other side of the antibody at the heavy-light chain interface like index and middle fingers (**Fig. 3C**). Analysis of the docked RBD-antibody complex showed that N477/K478 positioned at the junction formed by CDR H3, CDR L1 and L2 with slight clashes to a region centered at CDR H3 residue 100C (Kabat numbering) (**Fig. 3D**). Sequence alignment of CDR H3 of VH1-58-derived antibodies indicated that residue 100C varied in sidechain sizes, from serine in S2E12 to tyrosine in A23-58.1. Analysis showed that size of 100C reversely correlated with neutralization potency IC_80_ (*p*=0.046) (**Fig. 3D, Fig. 2C**), suggesting VH1-58 antibodies could alleviate escape imposed by the B.1.1.529 mutations through reduced side chain size at position 100C to minimize clashes from N477/K478.

**Figure 3.**
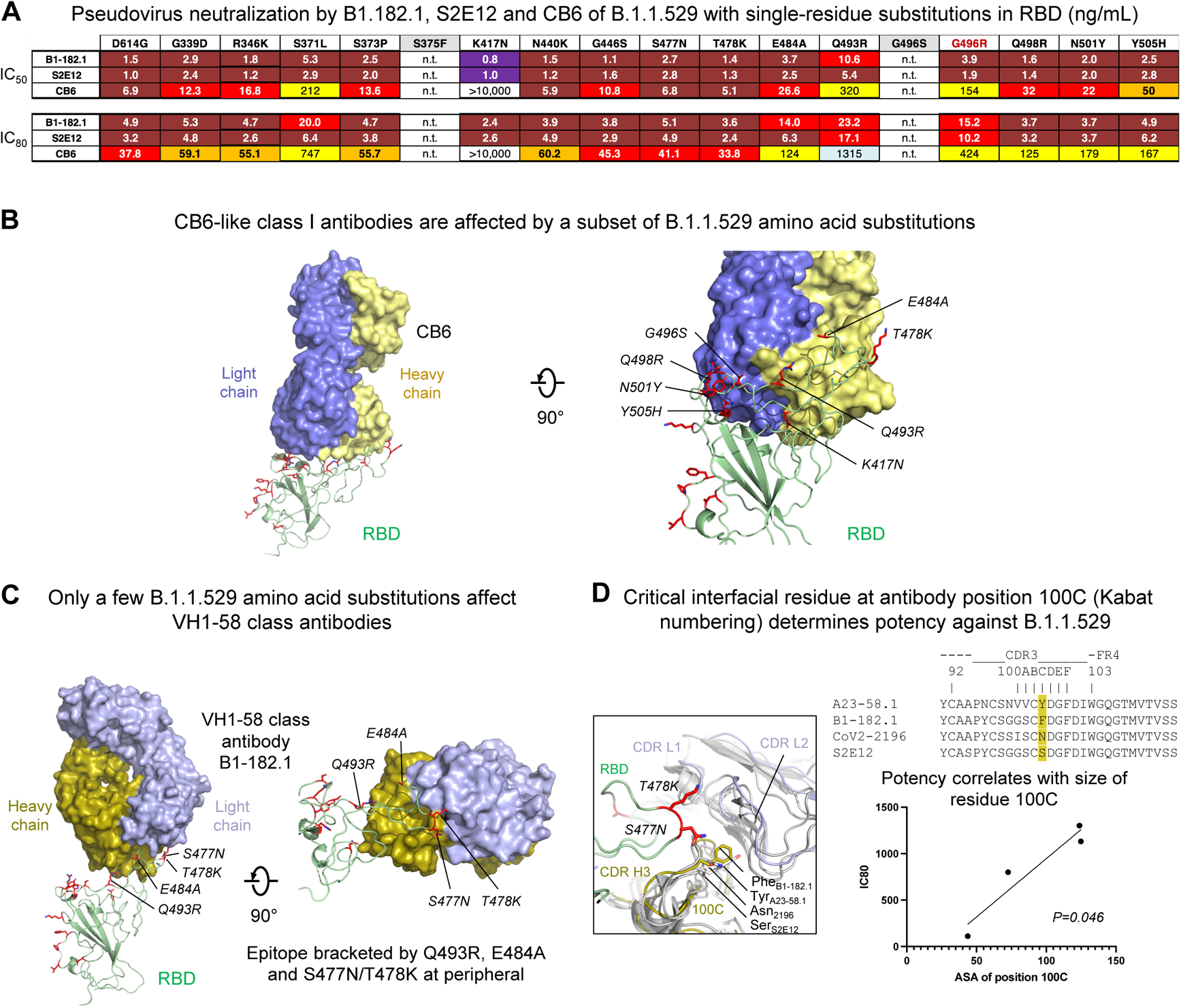
Functional and structural basis of Class I antibody neutralization and mechanistic basis of retained potency against B.1.1.529 VOC. A. Lentiviruses pseudotyped with SARS-CoV-2 spike proteins from D614G or D614G plus the indicated point mutations found within the B.1.1.529 spike were incubated with serial dilutions of the indicated antibodies, transduced 293T-ACE2 cells and IC_50_ and IC_80_ values determined. S375F and G496S viruses were not available and are shown as “not tested” (n.t.). G496R was available and substituted for G496S. Ranges are indicated with white (>10,000 ng/ml), light blue (>1000 to ≤10,000 ng/ml), yellow (>100 to ≤1000 ng/ml), orange (>50 to ≤100 ng/ml), red (>10 to ≤50 ng/ml), maroon (>1 to ≤10 ng/ml), and purple (≤1 ng/ml).
B. Mapping of B.1.1.529 amino acid substitutions at the epitope of Class I antibody CB6. RBD-bound CB6 was docked onto the B.1.1.529 spike structure. B.1.1.529 amino acid substitutions incompatible with CB6 binding were identified and labeled. K417N mutation caused clash in the center of the paratope. B.1.1.529 RBD is shown in green cartoon with amino acid substitutions in red sticks. CB6 is shown in surface representation with heavy and light chains colored yellow and slate, respectively.
C. Docking of RBD-bound VH1-58-derived Class I antibody B1-182.1 onto the B.1.1.529 spike structure identified 4 substitutions with potential steric hindrance. B1-182 is shown in surface representation with heavy and light chains colored olive and light blue. B.1.1.529 amino acid substitutions that may affect binding of VH1-58 antibodies were labeled.
D. Structural basis for effective neutralization of the B.1.1.529 VOC by VH1-58-derived antibodies. Even though VH1-58 antibodies, such as the S2E12, COV2-2196, A23-58.1 and B1-182.1, share high sequence homology (right, top), their neutralization potency against B.1.1.529 vary. Structural analysis indicated that CDR H3 residue 100C, located at the interfacial cavity formed by RBD, heavy and light chains, may determine their potency against B.1.1.529 (left). Size of this residue correlated with potency with two-tailed *p*=0.046 (right, bottom).

## Structural and functional basis of Class II antibody neutralization, escape and retained potency

We next sought to determine the functional basis of B.1.1.529 neutralization and escape for two Class II antibodies, LY-CoV555 (*26*) and A19-46.1 (*14*), which have B.1.1.529 IC_50_ of >10,000 and 223 ng/mL, respectively (**Fig 2C**). By assessing the impact of each of the single amino acid changes in RBD from B.1.1.529, we found that for LY-CoV555, either E484A or Q493R resulted in complete loss of LY-CoV555 neutralization (IC_50_ >10,000 ng/mL) (**Fig 4A**), while the same mutations did not affect A19-46.1. For A19-46.1, no individual mutation reduce neutralization to the level noted in B.1.1.529; S371L had the highest effect, reducing the IC_50_ to 72.3 ng/mL relative to 223 ng/mL for B.1.1.529. One potential explanation for this is that the Phe-Phe interaction, between 375 and 486, occurs in the context of three B1.1.529 alterations, S371L/S373P/S375F (**Fig. 1B**).

**Figure 4.**
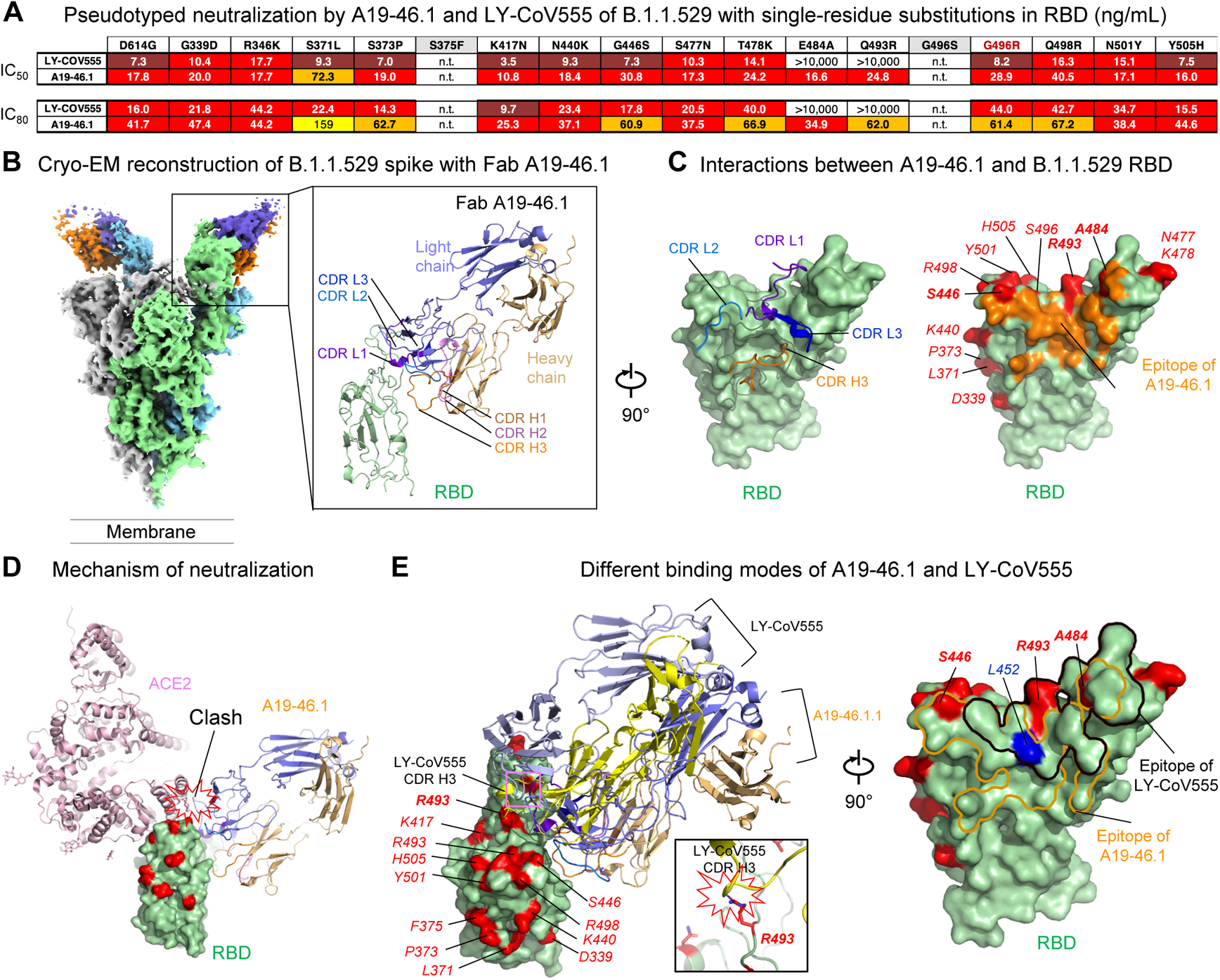
Functional and structural basis of Class II antibody binding, neutralization, and escape. A. Lentiviruses pseudotyped with SARS-CoV-2 spike proteins from D614G or D614G plus the indicated point mutations found within the B.1.1.529 spike were incubated with serial dilutions of the indicated antibodies, transduced 293T-ACE2 cells and IC_50_ and IC_80_ values determined. S375F and G496S viruses were not available and are shown as “not tested” (n.t.). G496R was available and substituted for G496S. Ranges are indicated with white (>10,000 ng/ml), light blue (>1000 to ≤10,000 ng/ml), yellow (>100 to ≤1000 ng/ml), orange (>50 to ≤100 ng/ml), red (>10 to ≤50 ng/ml), maroon (>1 to ≤10 ng/ml), and purple (≤1 ng/ml).
B. Cryo-EM structure of class II antibody A19-46.1 Fab in complex with the B.1.1.529 spike. Overall density map is shown to the left with protomers colored light green, gray and light cyan. Two A19-46.1 Fabs bound to the RBD in the up-conformation are shown in orange and slate. Structure of the RBD and A19-46.1 after local focused refinement was shown to the right in cartoon representation. The heavy chain CDRs are colored brown, pink and orange for CDR H1, CDR H2 and CDR H3, respectively. The light chain CDRs are colored marine purple blue, marine blue and blue for CDR L1, CDR L2 and CDR L3, respectively. The contour level of Cryo-EM map is 4.0 s.
C. Interaction between A19-46.1 and RBD. CDR H3 and all light chain CDRs were involved in binding of RBD (left). Epitope of A19-46.1 is shown in orange on the green B.1.1.529 RBD surface with amino acid substitutions colored red. Ser446, A484 and R493 are located at the edge of the epitope of Fab A19-46.1 (right). RBD residues are labeled with italicized font.
D. Binding of A19-46.1 to RDB prevents binding of the ACE2 receptor. ACE2 and A19-46.1 are shown in cartoon representation.
E. Comparison of binding modes to RBD for antibody A19-46.1 and LY-CoV555. Even though both antibodies target similar regions on RBD, different approaching angle caused clash between LY-CoV555 CDR H3 and B.1.1.529 mutation Arg493 (left and inset). B.1.1.529 mutations involved in binding of A19-46.1 are only at the edge of its epitope while both Arg493 and A484 locate in the middle of LY-CoV555 epitope (right). Leu452 to Arg mutation that knockouts A19-46.1 and LY-CoV555 binding in other SARS-CoV-2 variants is colored in blue.

To understand the structural basis of A19-46.1 neutralization of B.1.1.529, we obtained cryo-EM structure of the B.1.1.529 spike in complex with Fab A19-46.1 at 3.86 Å resolution (**Fig 4B, Fig. S5 and Table S2**). The structure revealed that two Fabs bound to the RBD in the “up” conformation in each spike with the third RBD in down position. Focused local refinement of the antibody-RBD region resolved the antibody-RBD interface (**Fig. 4B**, right). Consistent with previous mapping and negative stain EM data, A19-46.1 binds to a region on RBD generally targeted by the Class II antibodies with an angle approximately 45 degrees towards the viral membrane. Binding involves all light chain CDRs and only CDR H3 of the heavy chain and buried a total of 805 Å^2^ interface area from the antibody (**Fig. 4C**, left). With the light chain latching to the outer rim of the RBD and providing about 70 % of the binding surface, A19-46.1 uses its 17-residue-long CDR H3 to form parallel strand interactions with RBD residues 345-350 (**Fig. 4B, right**) like a sway brace. Docking RBD-bound ACE2 to the A19-46.1-RBD complex indicated that the bound antibody sterically clashes with ACE2 (**Fig. 4D**), providing the structural basis for its neutralization of B.1.1.529.

The 686 Å^2^ epitope of A19-46.1 is located within an RBD region that lacks amino acid changes found in B.1.1.529. Of the 15 amino acid changes on RBD, three of residues, S446, A484 and R493, positioned at the edge of epitope with their side chains contributing 8% of the binding surface. LY-CoV555, which targets the same region as a class II antibody, completely lost activity against B.1.1529. To gain structural insights on the viral escape of LY-CoV555, we superimposed the LY-CoV555-RBD complex onto the B.1.1.529 RBD. Even though LY-CoV555 approached the RBD with similar orientation to that of A19-46.1 (Fig. 4D), its epitope shifted up to the ridge of the RBD and embraced B.1.1.529 alterations A484 and R493 within the boundary (**Fig. 4D**). Inspection of the superimposed structures indicated B.1.1.529 alteration R493 caused steric clash with the CDR H3 of LY-CoV555, explaining the escape of B.1.1.529 from LY-CoV555 neutralization. Overall, the location of the epitope and the angle of approach allowed A19-46.1 to effectively neutralize B.1.1.529.

## Structural and functional basis of Class III antibody neutralization, escape and retained potency

To evaluate the functional basis of B.1.1.529 neutralization and escape for Class III antibodies and to understand how potent neutralization might be retained, we investigated a panel of Class III antibodies with differential potency, including A19-61.1, COV2-2130, S309 and LY-CoV1404 (**Fig 5A**). Assessment of the impact of each of the 15 mutations in RBD revealed the G446S amino acid change results in a complete loss in activity for A19-61.1; consistent with the complete loss of function of this antibody against B.1.1.529. For S309, we observed S373P and G496R to result in small changes to neutralization.

**Figure 5.**
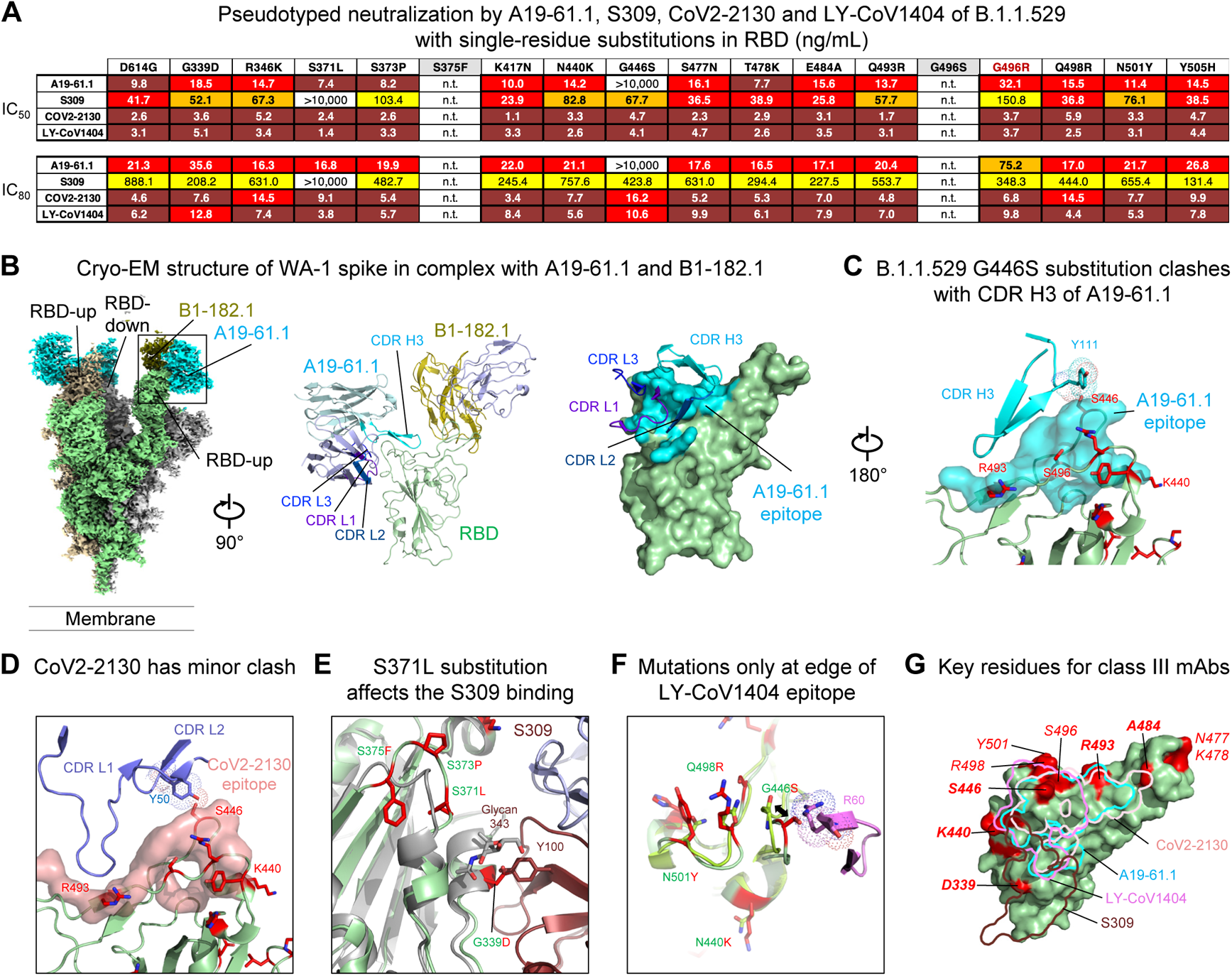
Functional and structural basis of Class III antibody binding, neutralization, and retained potency against the B.1.1.529 VOC. A. Lentiviruses pseudotyped with SARS-CoV-2 spike proteins from D614G or D614G plus the indicated point mutations found within the B.1.1.529 spike were incubated with serial dilutions of the indicated antibodies and IC_50_ and IC_80_ values determined. A19-61.1 and LY-COV1404 were assayed on 293T-ACE2 cells while S309 and CoV2-2130 were tested on 293 flpin-TMPRSS2-ACE2 cells. S375F and G496S viruses were not available and are shown as “not tested” (n.t.). G496R was available and substituted for G496S. Ranges are indicated with white (>10,000 ng/ml), light blue (>1000 to ≤10,000 ng/ml), yellow (>100 to ≤1000 ng/ml), orange (>50 to ≤100 ng/ml), red (>10 to ≤50 ng/ml), maroon (>1 to ≤10 ng/ml), and purple (≤1 ng/ml).
B. Cryo-EM structure of SARS-CoV-2 WA-1 spike in complex with class I antibody B1-182.1 and class III antibody A19-61.1 at 2.83 Å resolution. Overall density map is shown with protomers colored light green, gray and wheat. Two RBDs were in the up conformation with each binding both Fabs, and one RBD was in the down position with A19-61.1 bound. RBD, B1-182.1 and A19-61.1 are colored olive and cyan, respectively (left). Structure of the RBD with both Fabs bound after local focused refinement was shown to the right in cartoon representation. RBD is shown in green cartoon and antibody light chains are colored light blue (middle). Epitope of A19-61.1 is shown as cyan colored surface on RBD with interacting CDRs labeled (right). The contour level of cryo-EM map is 5.2s.
C. Structural basis of B.1.1.529 resistance to A19-61.1. Mapping of the A19-61.1 epitope onto the B.1.1.529 RBD indicated G446S clashed with CDR H3 of A19-61.1. RBD is shown in green cartoon with mutation residues in red sticks, epitope of A19-61.1 is shown in cyan surface.
D. Structural basis of CoV2-2130 neutralization of the B.1.1.529 VOC. Docking of the CoV2-2130 onto the B.1.1.529 RBD showed Y50 in CDR L2 posed a minor clash with S446. RBD is shown in green cartoon with mutation residues in red sticks, epitope of CoV2-2130 is shown in pink surface.
E. Structural basis of S309 neutralization of the B.1.1.529 VOC. Docked complex of S309 and B.1.1.529 RBD showed the S371L/S373P/S375F. Loop. Changed conformation, and the S371L mutation is adjacent to S309 epitope while G339D mutation located inside the epitope. D339 sidechain clash with CDR H3 Y100. B.1.1.529 RBD is shown in green cartoon with mutation residues in red sticks, WA-1 RBD is shown in gray cartoon.
F. Structural basis of LY-CoV1404 neutralization of the B.1.1.529 VOC. Docking of the LY-CoV1404 onto the B.1.1.529 RBD identified 4 amino acid substitutions in the epitope with G446S causing potential clash with CDR H2 R60. However, comparison of both LY-CoV1404-bound and non-bound B.1.1.529 RBD indicated the S446 loop has the flexibility to allow LY-CoV1404 binding. B.1.1.529 residues at LY-CoV1404 epitope are shown as red sticks with corresponding WA-1 residues as green. sticks. CDR H3 is shown in cartoon representation and colored magenta.
G. Overlay of epitope footprints of class III antibodies onto the B.1.1.529 RBD. B.1.1.529 RBD amino acid substitution locations are colored red on green surface.

Surprisingly, while S309 retains moderate neutralizing activity against B.1.1.529, we found the S371L amino acid change to abolish S309 neutralization. This suggests that combinations of S371L with other B.1.1.529 mutations can result in structural changes in spike that allows S309 to partially overcome the S371L change. The evaluation of COV2-2130 did not identify significant differences in neutralization, suggesting a role for an untested mutation or combinations of amino acid changes for the decrease in neutralization potency observed against the full virus. Finally, consistent with the overall high potency of LY-CoV1404 against all tested VOCs, we did not identify an amino acid change that impacted its function. To understand the structural basis of Class III antibody neutralization and viral escape, we determined the cryo-EM structure of WA-1 S2P in complex with Fab A19-61.1 (and Fab B1-182.1 to aid EM resolution of local refinement) at 2.83 Å resolution (**Fig. 5B, Fig. S6 and Table S1**). The structure revealed that two RBDs were in the up-conformation with both antibodies bound, and the third RBD was in the down-position with only A19-61.1 bound, indicating A19-61.1 could recognize RBD in both up and down conformation (**Fig. 5B**). Local refinement of the RBD-Fab A19-61.1 region showed that A19-61.1 targeted the Class III epitope with interactions provided by the 18-residue-long CDR H3 from the heavy chain, and all CDRs from the light chain (**Fig. 5B**). Docking the A19-61.1 structure to the B.1.1.529 spike structure indicated B.1.1.529 mutations S446, R493 and S496 might interfere with A19-61.1. Analysis of the side chain interaction identified Y111 in CDR H3 posed severe clash with S446 in RBD that could not be resolved by loop flexibility (**Fig. 5C**), explaining the loss of A19-61.1 neutralization against G446S-containding SARS-CoV-2 variants.

Neutralization assays indicated that COV2-2130, S309 and LY-CoV1404 retained neutralization potency against B.1.1.529. Docking of CoV2-2130 indicated that CoV2-2130 targeted a very similar epitope to that of A19-61.1 with interactions mainly mediated by its CDR L1 and L2 and avoiding close contact with R493 and S496. However, the OH group at the tip of Y50 in CDR L2 posed a minor clash with S446 in RBD, explaining the structural basis for the partial conservation of neutralization by CoV2-2130 (**Fig. 5D**). Antibody S309 showed higher potency against B.1.1.529 than CoV2-2130. Docked complex of S309 and RBD showed the G339D mutation is located inside the epitope and clashes with CDR H3 Y100, however, the void space between S309 and RBD might accommodate an alternate tyrosine rotamer. The S371L/S373P/S375F mutations changed the conformation of their residing loop and may push the glycan on N343 towards S309 to reduce binding (**Fig. 5E**). LY-CoV1404 was not affected by B.1.1.529 mutations.

Docking of the LY-CoV1404 onto the B.1.1.529 RBD identified four amino acid substitutions located at the edge of its epitope. Three of the residues, K440, R498 and Y501, only make limited side chain interactions with LY-CoV1404. The 4^th^ residue, G446S, appeared to cause a potential clash with CDR H2 R60. However, comparison of both LY-CoV1404-bound and non-bound RBD indicated the loop where S446 resided had conformational flexibility to allow LY-CoV1404 binding (**Fig. 5F**). Overall, the epitopes to Class III antibodies were mainly located on mutation-free RDB surfaces with edges contacting a few B.1.1.529 alterations (**Fig. 5G**). LY-CoV1404 retained high potency by accommodating all four B.1.1.529 alteration at edge of its epitope by exploiting loop mobility or by minimizing side chain interactions.

## Synergistic neutralization by the combination of B1-182-1 and A19-46.1

We previously reported that the combination of B1-182.1 and either A19-46.1 or A19-61.1 mitigated mutational escape in an *in vitro* virus escape assay (*14*); suggesting the possibility of synergistic neutralization. We therefore hypothesized that there are combinations of antibodies which lead to an increase in B1.1.529 neutralization beyond that of the two antibodies alone. As a test of this hypothesis, we determined the neutralization of B.1.1.529 pseudotyped viruses by clinically utilized cocktails or various combinations of B1-182.1, A19-46.1, A19-61.1, LY-CoV1404, ADG2 and S309. Of the 10 combinations evaluated only COV2-2196/COV2-2130, B1-182.1/A19-46.1 and B1-182.1/S309 neutralized B.1.1.529 with an appreciably improved potency (*i.e.,* IC_50_ of 50.8, 28.3 and 58.1 ng/mL) over the individual component antibodies (**Fig 6A,B**). Each of these utilized a VH-158 supersite antibody and showed a 5 to 115-fold improvement over the component antibodies (**Fig 6B**), suggesting an effect that is more than an additive for the specific combination against B.1.1.529.

**Figure 6.**
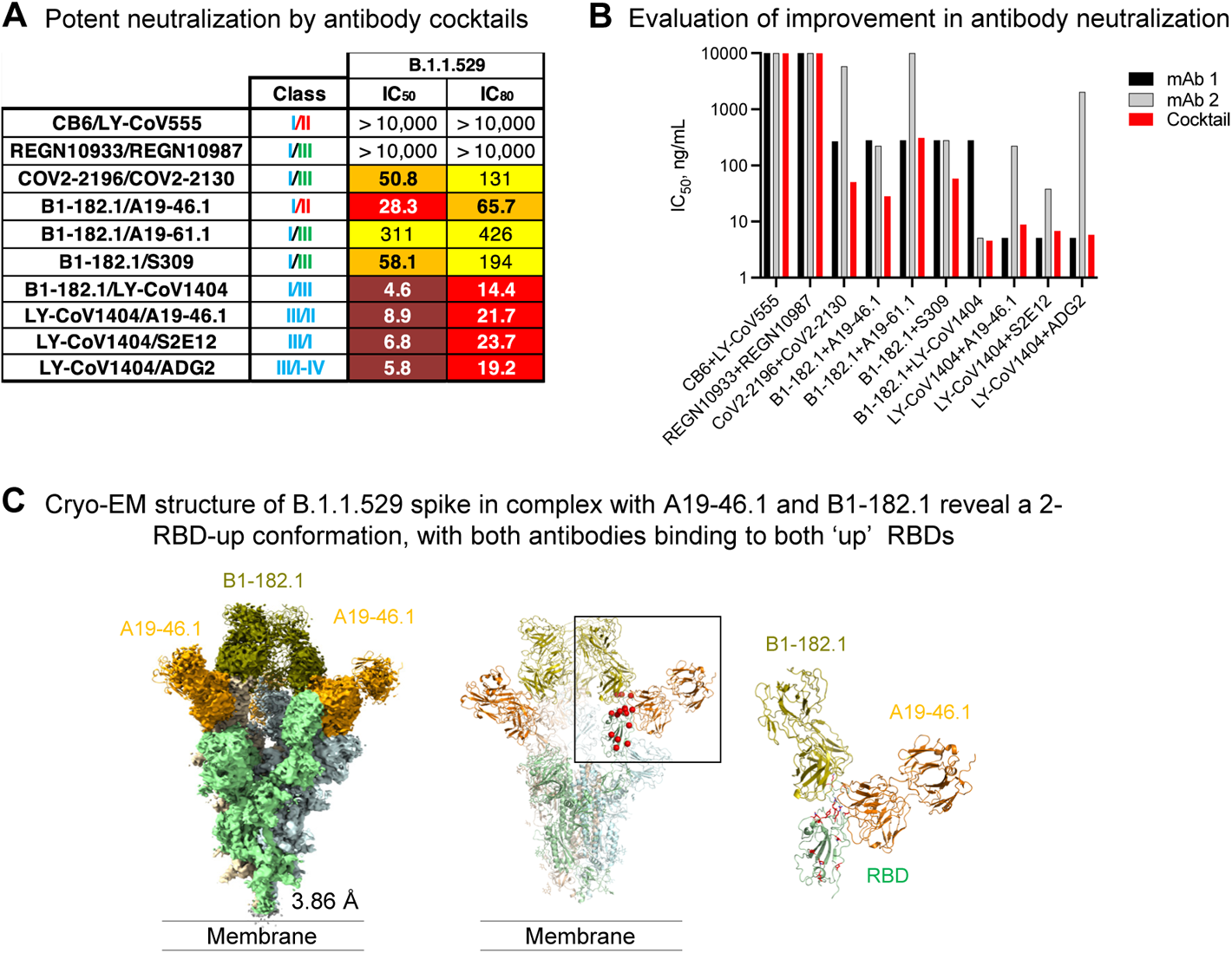
Potent neutralization of SARS-CoV-2 B.1.1.529 using combinations of antibodies. A. Lentiviruses pseudotyped with SARS-CoV-2 spike proteins from D614G or D614G plus the indicated point mutations found within the B.1.1.529 spike were incubated with serial dilutions of the indicated combination of antibodies and IC_50_ and IC_80_ values determined. S375F and G496S viruses were not available and are shown as “not tested” (n.t.). G496R was available and substituted for G496S. Ranges are indicated with white (>10,000 ng/ml), light blue (>1000 to ≤10,000 ng/ml), yellow (>100 to ≤1000 ng/ml), orange (>50 to ≤100 ng/ml), red (>10 to ≤50 ng/ml), maroon (>1 to ≤10 ng/ml), and purple (≤1 ng/ml).
B. Neutralization IC_50_ (ng/mL) values for each of the indicated cocktail (x-axis) or its component antibodies. The IC_50_ for first antibody is listed as mAb1 (black), the second antibody as mAb2 (grey) or cocktail (red).
C. Cryo-EM structure of B.1.1.529 spike in complex with antibodies A19-46.1 and B-182.1 at 3.86 Å resolution. Overall density map is shown to the left with protomers colored light green, wheat and light cyan (left). All RBD are in up-conformation with both Fabs bound (middle). Binding of one Fab (such as B1-182.1) induces RBD into the up-conformation and potentially facilitates binding of the other Fab (such as A19-46.1) which only recognizes the up-conformation of RBD (right). A19-46.1 and B-182.1 are shown in orange and olive color, respectively. The contour level of cryo-EM map is 6.5 s.

To understand the structural basis of the improved neutralization by the cocktail of B1-182.1 and A19-46.1 we determined the cryo-EM structure of the B.1.1.529 S2P spike in complex with Fabs of B1-182.1 and A19-46.1 at 3.86 Å resolution (**Fig. 6C, Fig. S7 and Table S1**). The prevalent 3D reconstruction revealed that the spike recognized by the combination of these two antibodies was the 3-RBD-up conformation with both Fabs bound to each RBD (Fabs on one of the RBDs were lower in occupancy). The spike had a 1.6 Å RMSD relative to the 3-RBD-up WA-1 structure (PDB ID: 7KMS). Overall, the structure showed that both Class I and II antibodies were capable of simultaneously recognizing the same RBD, and the combination increased the overall stoichiometry compared to two Fabs per trimer observed in the S2P-A19-46.1 structure described above. Of all the antibodies tested, we note that all VH1-58-derived antibodies retained reasonable level of neutralization against B.1.1.529 while members of other antibody classes suffered complete loss of activity. VH1-58 antibodies have minimal numbers of impacting B.1.1.529 alterations in their epitopes and can evolve means to alleviate the impact. We speculate that binding of the first antibody induced the spike into RBD-up-conformation and facilitated binding of the second RBD-up-conformation preferring antibody, thereby synergistically increasing the neutralization potency of the cocktail compared to the individual antibodies.

## Discussion

SARS-CoV2 variants of concern provide a window into the co-evolution of key host-pathogen interactions between the viral spike, human ACE2 receptor and the human immune system. The RBD is a major target for neutralizing antibodies in both survivors and vaccinees. Since 15 of the 37 mutations in the B.1.1.529 variant spike reside within the RBD, there is a great need to understand the mechanisms by which RBD variations evolve, what constraints exist on the evolution and whether there are approaches that can be taken to exploit this understanding to develop and maintain effective antibody therapeutics and vaccines. The B.1.1.529 cryo-EM structure showed a cluster of RBD mutations proximal to the ACE2-binding surface, which alters the electrostatic potential of the interface. However, in contrast to studies that showed increased human ACE2 affinity to mutated and variant RBD subdomains (*20, 34, 35*), our studies found that in the context of trimeric spike proteins, variant amino acid changes did not provide a biologically meaningful alteration in affinity. This suggests that there is either no further fitness benefit to be gained by improving affinity, that affinity improving changes are being used to compensate for mutations that are deleterious for ACE2 binding but allow immune escape, or both.

We used a series of functional and structural studies to define the mechanisms by which B.1.1.529 is either neutralized by or mediates escape from host immunity. To functionally frame our analyses, we utilized the Barnes classification, which categorizes antibodies based on their binding to the ACE2 binding site and the position of RBD. Our findings for Class I VH1-58 supersite showed that B.1.1.529 requires a series of mutation that are not individually deleterious to bracket the antibody and reduce its potency. Notably our data suggests that VH1-58 antibodies can alleviate the deleterious impact of this pinching effect by reducing the size of CDR H3 residue 100C to avoid clashes from B.1.1.529 mutations. Since VH1-58 supersite are amongst the most potent and broadly neutralizing anti-SARS-CoV-2 antibodies (*14, 25, 29, 36*), our findings point the way toward structure-based designs of existing antibodies to mitigate against amino acid changes at these positions.

For the Class II antibody A19-46.1, the angle of approach and a long-CDRH3 combine allow it to target the mutation-free face on RBD and minimize contacting the mutations on the ridge of B.1.1.529 RBD. We observed that A19-46.1 binding requires the RBD-up conformation, and that the S371L substitution, which is located away from the A19-46.1 epitope and near the RBD hinge, partially reduces the neutralization of A19-46.1. Comparing the effect of S371L on neutralization by A19-46.1 and LY-CoV555 (**Fig. 4A**), which recognizes both RBD-up and -down conformation, suggested that L371 (and potentially P373/F375) is critical for controlling the RBD-up or -down conformation in B.1.1.529. This concept is supported by the finding that combination with a Class I antibody (such as B1-182.1) synergistically enhances A19-46.1 neutralization (**Fig. 6A**).

For Class III antibodies, only one prototype antibody showed complete loss of B.1.1.529 neutralization. Using structural and functional approaches we determined that viral escape was mediated by the G446S amino acid change. This result indicates that potent Class III antibodies might be induced through structure-based vaccine designs that mask residue 446 in RBD. Additionally, the existence of G446S sensitive and resistant antibodies with significant epitope overlap suggest the use of spikes with G446S substitution can be utilized to evaluate the quality of Class III immune response in serum-based epitope mapping assays (*37, 38*).

Our analysis of antibodies of clinical importance is consistent with previous reports (*32, 39–41*) and showed that S309 and COV2-2196 neutralized to similar degrees. Importantly, we report that unlike other antibodies, the highly potent LY-CoV1404 does not lose neutralization potency against B.1.1.529. We identified combinations of antibodies that show more than additive increases in neutralization against B.1.1.529, including COV2-2196/COV2-2130, B1-182.1/A19-46.1 and B1-182.1/S309. Each pair contains a VH1-58 supersite antibody that binds RBD in the up position. We speculated that pairing antibodies that neutralize better in the up-RBD conformation with these VH1-58 antibodies may provide a mechanism for better neutralization by the former. The S371L/S373P/S375F alterations in the RBD-up protomer form interprotomer interactions to RBD in the RBD-down protomer and stabilize the B.1.1.529 spike into a single-RBD-up conformation. RBD-up-preferring antibody like the VH1-58-derived B1-182.1, which is not affected by S371L substitution, can effectively break up the interaction to induce the 3-RBD-up conformation and therefore, enhance binding of other antibodies (such as A19-46.1) that require the RBD up-conformation. The identification of SARS-CoV-2 monoclonal antibodies that cooperatively function is similar to that seen previously for other viruses (*42*), and supports the concept of using combinations to both enhance potency and mitigate the risk of escape.

## Acknowledgments

We thank N.A. Doria-Rose, W.-P. Kong, S. O’Dell and S.D. Schmitt and for assistance in B.1.1.529 plasmid production and distribution, M. Kanekiyo for cell line assistance, J. Stuckey and S. Wang for assistance with manuscript preparation and submission, and members of the Virology Laboratory, Vaccine Research Center, for discussions and comments on the manuscript. We thank S Žentelis, E. Lameignere and K. Westerndorf for antibody LY-CoV1404. We are grateful to T. Edwards and T.L. Fox of NCEF for cryo-EM data collection and for technical assistance with cryo-EM data processing.

## Author contributions

T.Z., L.W., J.M. and A.P. designed experiments and analyzed data. L.W., A.P., Y.Z., D.R.H., C.A.T., A.S.O., E.S.Y., M.C. (Man Chen), K.L. and E.S.D.S. performed experiments. L.W., J.M., A.S.O., W.S., M.C. (Misook Choe), I-T.T., A.C., T.L., B.Z. produced proteins, antibodies and other reagents. T.Z. led electron microscopy studies assisted by C.J., T.S. and Y.T.. J.M., B.S.G., J.R.M., N.J.S. and P.D.K. supervised experiments. T.Z., L.W., J.M., N.J.S., and P.D.K wrote the manuscript with help from all authors.

## Data and materials availability

All data is available in the main text or the supplementary materials. Atomic coordinates and cryo-EM maps of the reported structure have been deposited into the Protein Data Bank and Electron Microscopy Data Bank under the session codes PDB 7TB4 and EMD-25792 for the SARS-CoV-2 B.1.1.529 VOC spike, PDB 7TCA and EMD-25807 for SARS-CoV-2 B.1.1.529 VOC spike in complex with antibody A19-46.1. PDB 7TC9 and EMD-25806 for local refinement of the SARS-CoV-2 B.1.1.529 VOC RBD in complex with antibody A19-46.1, PDB 7TCC and EMD-25808 for SARS-CoV-2 B.1.1.529 VOC spike in complex with antibodies B1-182.1 and A19-46.1. PDB 7TB8 and EMD-25794 for SARS-CoV-2 WA-1 spike in complex with antibodies A19-61.1 and B1-182.1, PDB 7TBF and EMD-25797 for local refinement of the SARS-CoV-2 WA-1 RBD in complex with antibodies A19-61.1 and antibody B1-182.1. Original materials in this manuscript are available under a materials transfer agreement with the National Institutes of Health.

## Supplementary Materials

Materials and Methods

Figures S1-S7

Tables S1-S2

## Supplementary Materials

### Materials and Methods

#### Expression and purification of proteins

Soluble 2P-stabilized SARS-CoV-2 spike proteins were expressed by transient transfection(*6, 43*). Briefly, plasmid was transfected using Expifectamine into Expi293 cells (Life Technology) and the cultures enhanced 16-24 hours post-transfection. Following 4-5 days incubations at 120 rpm, 37 °C, 9% CO2, supernatant was harvested, clarified via centrifugation, and buffer exchanged into 1X PBS. Protein of interests were then isolated by affinity chromatography using Ni-NTA resin (Roche) followed by size exclusion chromatography on a Superose 6 increase 10/300 column (GE healthcare).

Expression and purification of biotinylated S2P used in binding studies were produced by an in-column biotinylation method as previously described (*43*). Using full-length SARS-Cov2 S and human ACE2 cDNA ORF clone vector (Sino Biological, Inc) as the template to the ACE2 dimer proteins. The ACE2 PCR fragment (1~740aa) was digested with Xbal and BamHI and cloned into the VRC8400 with Avi-HRV3C-single chain-human Fc-his (6x) tag on the C-terminal. All constructs were confirmed by sequencing. Proteins were expressed in Expi293 cells by transfection with expression vectors encoding corresponding genes. The transfected cells were cultured in shaker incubator at 120 rpm, 37 °C, 9% CO2 for 4~5 days. Culture supernatants were harvested and filtered, and proteins were purified through a Hispur Ni-NTA resin (Thermo Scientific) and following a Hiload 16/600 Superdex 200 column (GE healthcare, Piscataway NJ) according to manufacturer’s instructions. The protein purity was confirmed by SDS-PAGE.

#### Synthesis, cloning and expression of monoclonal antibodies

Sequences were selected for synthesis to sample expanded clonal lineages within our dataset and convergent rearrangements both among donors in our cohort and compared to the public literature. In addition, we synthesized a variety of sequences designed to be representative of the whole dataset along several dimensions, including apparent epitope based on flow data; V gene usage; somatic hypermutation levels; CDRH3 length; and isotype. Variable heavy chain sequences were human codon optimized, synthesized and cloned into a VRC8400 (CMV/R expression vector)-based IgG1 vector containing an HRV3C protease site (*44*) as previously described (*45*). Similarly, variable lambda and kappa light chain sequences were human codon optimized, synthesized and cloned into CMV/R-based lambda or kappa chain expression vectors, as appropriate (Genscript). ADG2 was kindly provided by Dr. Laura M Walker (Adagio Therapeutics, Inc., Waltham, MA) (*23*) and LY-CoV1404 by Dr. Stefanie Žentelis, Dr Emilie Lameignere and Kathryn Westendorf, MSc (AbCellera, Inc., Canada) (*24*). Previously published antibody vectors for LY-COV555 were used (*26*). For antibodies where vectors were unavailable (e.g., S309, CB6, REGN10933, REGN10987, COV2-2196, COV2-2130, CT-P59, C144, C135, S2E12) (*12, 13, 21, 22, 25, 27–31*), published amino acids sequences were used for synthesis and cloning into corresponding pVRC8400 vectors (Genscript) (*46, 47*). For antibody expression, equal amounts of heavy and light chain plasmid DNA were transfected into Expi293 cells (Life Technology) by using Expi293 transfection reagent (Life Technology). The transfected cells were cultured in shaker incubator at 120 rpm, 37 °C, 9% CO2 for 4~5 days. Culture supernatants were harvested and filtered, mAbs were purified over Protein A (GE Health Science) columns. Each antibody was eluted with IgG elution buffer (Pierce) and immediately neutralized with one tenth volume of 1M Tris-HCL pH 8.0. The antibodies were then buffer exchanged as least twice in PBS by dialysis.

#### Full-length S constructs

Codon optimized cDNAs encoding full-length S from SARS CoV-2 (GenBank ID: QHD43416.1) were synthesized, cloned into the mammalian expression vector VRC8400 (*46, 47*) and confirmed by sequencing. S containing D614G amino acid change was generated using the wt S sequence. Other variants containing single or multiple aa changes in the S gene from the S wt or D614G were made by mutagenesis using QuickChange lightning Multi Site-Directed Mutagenesis Kit (cat # 210515, Agilent) or via synthesis and cloning (Genscript). The S variants tested are B.1.351 (L18F, D80A, D215G, (L242-244)del, R246I, K417N, E484K, N501Y, A701V), P.1 (L18F, T20N, P26S, D138Y, R190S, K417T, E484K, N501Y, D614G, H655Y, T1027I, V1176F), B.1.1.7 (H69del, V70del, Y144del, N501Y, A570D, D614G, P681H, T716I, S982A, D1118H), B.1.617.2 (T19R, G142D, E156del, F157del, R158G, L452R, T478K, D614G, P681R, D950N), B.1.1.529 (A67V, H69del, V70del, T95I, G142D, V143del, Y144del, Y145del, N211del, L212I, ins214EPE, G339D, S371L, S373P, S375F, K417N, N440K, G446S, S477N, T478K, E484A, Q493R, G496S, Q498R, N501Y, Y505H, T547K, D614G, H655Y, N679K, P681H, N764K, D796Y, N856K, Q954H, N969K, L981F). The S genes containing single RBD amino acid changes from the B.1.1.529 variant were generated based on D614G construct by mutagenesis. These full-length S plasmids were used for pseudovirus production and for cell surface binding assays.

#### Generation of 293 Flpin-TMPRSS2-ACE2 cell line

293 Flpin-TMPRSS2-ACE2 isogenic cell line was prepared by co-transfecting pCDNA5/FRT plasmid encoding TMPRSS2-T2A-ACE2 and pOG44 plasmid encoding Flp recombinase in 293 Flpin parental cell line (Thermo Fisher, Cat R75007). Cells expressing TMPRSS2-ACE2 were selected using Hygromycin at 100 micrograms/ml. TMPRSS2 and ACE2 expression profiles in 293 Flpin-TMPSS2-ACE2 were characterized by flow cytometry using a mouse monoclonal antibody against TMPRSS2 (MillliporeSigma) followed by an anti-mouse IgG1 APC conjugate (Jackson laboratories) and a molecular probe containing the SARS-CoV-2 receptor binding domain tagged with biotin (Sino biological) followed by staining with a BV421 conjugated streptavidin probe (BD Biosciences).

#### Pseudovirus neutralization assay

S-containing lentiviral pseudovirions were produced by co-transfection of packaging plasmid pCMVdR8.2, transducing plasmid pHR’ CMV-Luc, a TMPRSS2 plasmid and S plasmids from SARS CoV-2 variants into 293T cells using Lipofectamine 3000 transfection reagent (L3000-001, ThermoFisher Scientific, Asheville, NC) (*48, 49*). 293T-ACE2 cells (provided by Dr. Michael Farzan) or 293 flpin-TMPRSS2-ACE2 cells were plated into 96-well white/black Isoplates (PerkinElmer, Waltham, MA) at 75,00 cells per well the day before infection of SARS CoV-2 pseudovirus. Serial dilutions of mAbs were mixed with titrated pseudovirus, incubated for 45 minutes at 37°C and added to cells in triplicate. 293 flpin-TMPRSS2-ACE2 cells were used for some of Class III antibodies like S309 and COV2-2130 while 293T-ACE2 cells were used for the rest of antibodies. Following 2 h of incubation, wells were replenished with 150 ml of fresh media. Cells were lysed 72 h later, and luciferase activity was measured with Microbeta (Perking Elmer). Percent neutralization and neutralization IC50s, IC80s were calculated using GraphPad Prism 8.0.2.

#### Cell surface binding

HEK293T cells were transiently transfected with plasmids encoding full length SARS CoV-2 spike variants using lipofectamine 3000 (L3000-001, ThermoFisher) following manufacturer’s protocol. After 40 hours, the cells were harvested and incubated with monoclonal antibodies (0.5 μg/ml) or biotinylated-human ACE2 (10 μg/ml, AC2-H82F9, Acro Biosystems) for 30 minutes. After incubation with the antibodies or ACE2, the cells were washed and incubated with an allophycocyanin conjugated anti-human IgG (709-136-149, Jackson Immunoresearch Laboratories) or BV421 conjugated streptavidin conjugate for another 30 minutes. The cells were then washed and fixed with 1% paraformaldehyde (15712-S, Electron Microscopy Sciences). The samples were then acquired in a BD LSRFortessa X-50 flow cytometer (BD biosciences) and analyzed using Flowjo (BD biosciences). Mean fluorescent intensity (MFI) for antibody binding to S D614G was set up as 100%. The MFI of ACE2 of antibody binding to each variant was normalized to S D614G.

#### Production of Fab fragments from monoclonal antibodies

To generate mAb-Fab, IgG was incubated with HRV3C protease (EMD Millipore) at a ratio of 100 units per 10 mg IgG with HRV 3C Protease Cleavage Buffer (150 mM NaCl, 50 mM Tris-HCl, pH 7.5) at 4°C overnight. Fab was purified by collecting flowthrough from Protein A column (GE Health Science), and Fab purity was confirmed by SDS-PAGE.

#### Determination of binding kinetics of ACE2

Binding kinetics and affinities of ACE2 to SARS-CoV-2 S2P variants were assessed by surface plasma resonance on a Biacore S-200 (GE Healthcare) at 25°C in the HBS-EP+ buffer (10 mM HEPES, pH 7.4, 150 mM NaCl, 3 mM EDTA, and 0.05% surfactant P20). Fc-reactive anti-human IgG antibody (Cytiva) wa coupled to the CM5 chip to approximately 10,000 RU, and dimeric, Fc-tagged ACE2 (ACRO Biosystems) at 35 μg/mL was captured for 60 seconds at 10 μL/min to a response of approximately 200 RU. Serially diluted SARS-CoV-2 S2P variants starting at 100 nM were flowed through the sample and reference channels for 180 seconds at 30 μL/min, followed by a 300 second dissociation phase at 30 uL/min. The chip was regenerated using 3 M MgCl_2_ for 30 seconds at 50 μL/min. Blank sensorgrams were obtained with HBS-EP+ buffer. Blank-corrected sensorgrams of the S2P concentration series were fitted globally with Biacore S200 evaluation software using a 1:1 model of binding. Plots were generated using GraphPad Prism.

#### Cryo-EM specimen preparation and data collection

Cryo-EM grids for the B.1.1.529 spike stabilized with the “2P” mutations were prepared at 0.5 mg/ml in a buffer containing 10 mM HEPES, pH 7.5 and 150 mM NaCl. For the spike-Fab complexes, the stabilized SARS-CoV-2 spikes of B.1.1.529 or WA-1were1. were mixed with Fab or Fab combinations at a molar ratio of 1.2 Fab per protomer in PBS with final spike protein concentration at 0.5 mg/ml. n-Dodecyl β-D-maltoside (DDM) detergent was added to the protein complex mixtures shortly before vitrification to a concentration of 0.005%. Quantifoil R 2/2 gold grids were subjected to glow discharging in a PELCO easiGlow device (air pressure: 0.39 mBar, current: 20 mA, duration: 30 s) immediately before specimen preparation. Cryo-EM grids were prepared using an FEI Vitrobot Mark IV plunger with the following settings: chamber temperature of 4°C, chamber humidity of 95%, blotting force of −5, blotting time of 2 to 3.5 s, and drop volume of 2.7 µl. Datasets were collected at the National CryoEM Facility (NCEF), National Cancer Institute, on a Thermo Scientific Titan Krios G3 electron microscope equipped with a Gatan Quantum GIF energy filter (slit width: 20 eV) and a Gatan K3 direct electron detector (Table S2). Four movies per hole were recorded in the counting mode using Latitude software. The dose rate was 14.65 e-/s/pixel.

#### Cryo-EM data processing and model fitting

Data process workflow, including motion correction, CTF estimation, particle picking and extraction, 2D classification, ab initio reconstruction, homogeneous refinement, heterogeneous refinement, non-uniform refinement, local refinement and local resolution estimation, were carried out with C1 symmetry in cryoSPARC 3.3 (*50*). For local refinement to resolve the RBD-antibody interface, a mask for the entire spike-antibody complex without the RBD-antibody region was used to extract the particles and a mask encompassing the RBD-antibody region was used for refinement. The overall resolution was 3.29 Å for the map of B.1.1.529 spike alone structure, 3.85 Å for the map of B.1.1.529 spike in complex with A19-46.1, 2.83 Å for the map of WA-1 spike in complex with A19-61.1 and B1-182.1, and 3..86Å for the map of B.1.1.529 spike in complex with A19-46.1 and B1-182.1. The coordinates for the SARS-CoV-2 spike with B1-182.1 molecules bound at pH 7.4 (PDB ID: 7MM0) were used as initial models for fitting the cryo-EM map. Iterative manual model building and real-space refinement were carried out in Coot (48) and in Phenix (*51*), respectively. Molprobity (*52*) was used to validate geometry and check structure quality at each iteration step. UCSF Chimera and ChimeraX were used for map fitting and manipulation (*53*).Data process workflow, including motion correction, CTF estimation, particle picking and extraction, 2D classification, ab initio reconstruction, homogeneous refinement, heterogeneous refinement, non-uniform refinement, local refinement and local resolution estimation, were carried out with C1 symmetry in cryoSPARC 3.3 (*50*). For local refinement to resolve the RBD-antibody interface, a mask for the entire spike-antibody complex without the RBD-antibody region was used to extract the particles and a mask encompassing the RBD-antibody region was used for refinement. The overall resolution was 3.29 Å for the map of B.1.1.529 spike alone structure, 3.85 Å for the map of B.1.1.529 spike in complex with A19-46.1, 2.83 Å for the map of WA-1 spike in complex with A19-61.1 and B1-182.1, and 3..86Å for the map of B.1.1.529 spike in complex with A19-46.1 and B1-182.1. The coordinates for the SARS-CoV-2 spike with B1-182.1 molecules bound at pH 7.4 (PDB ID: 7MM0) were used as initial models for fitting the cryo-EM map. Iterative manual model building and real-space refinement were carried out in Coot (48) and in Phenix 1.19.2 (*51*), respectively. Molprobity (*52*) was used to validate geometry and check structure quality at each iteration step. Map fitting and manipulation and display were performed with UCSF Chimera and ChimeraX were used for map fitting and manipulation (*53*).

#### Differential Scanning Calorimetry (DSC)

DSC measurements were performed using a VP-ITC (Microcal) instrument. Spike samples were diluted to 0.125mg/ml in PBS and scanned from 20 to 95°C at a rate of 1°C per minute. Thermal denaturation (Tm) temperature and total enthalpy of unfolding was calculated using the Microcal analysis system in Origin.

#### Biolayer interferometry binding assay

The antibody binding panel was performed on a FortéBio Octet HTX instrument with black, tilted 384-well plates (Greiner Bio-One). All steps of pre-soaking, binding and dissociation were performed in PBS with 1% BSA at pH 7.4. IgGs and dACE2-Fc were loaded onto Anti-Human Fc Sensor Tips (FortéBio) at a concentration of 1-4μg/mL, resulting in a load response of 0.85-1.5 nm. The plates were agitated at 1,000 rpm and the experiment run at 30°C. Antibodies and ACE2 were loaded onto the tips for 2 minutes, bound to 100nM S2P protein for 5 minutes and dissociated in buffer for 5 minutes. Reference well subtraction was performed with the Data Analysis Software HT v12.0 (FortéBio). The graphs were generated in GraphPad Prism.

---

**Figure S1.**
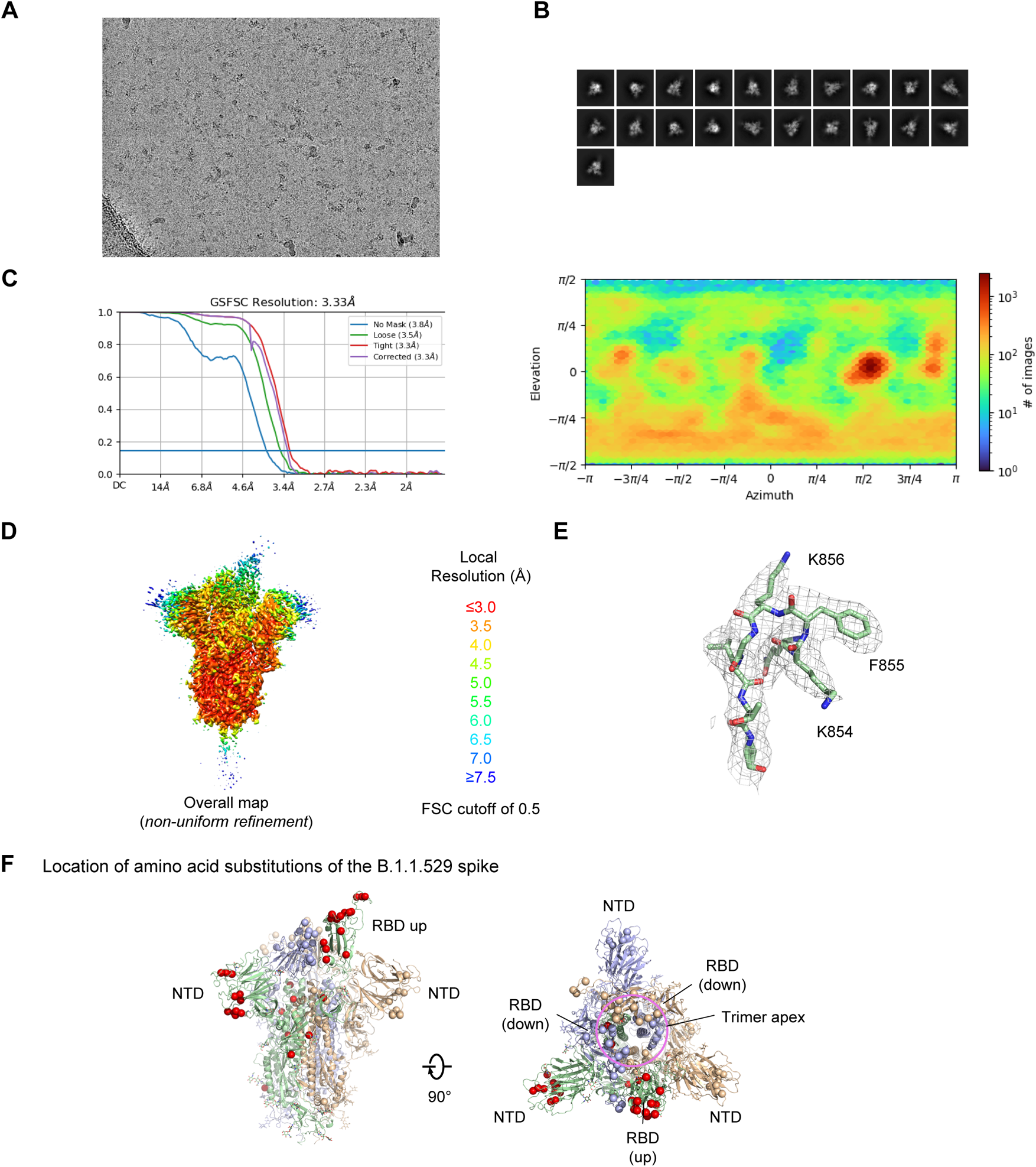
Cryo-EM details of the SARS-CoV-2 B.1.1.529 spike. A. Representative micrograph.
B. Representative 2D class averages are shown.
C. The gold-standard Fourier shell correlation resulted in a resolution of 3.33 Å for the overall map using non-uniform refinement with C1 symmetry (left panel); the orientations of all particles used in the final refinement are shown as a heatmap (right panel).
D. The local resolution of the final overall map is shown contoured at 0.373 (5.7σ). Resolution estimation was generated through cryoSPARC using an FSC cutoff of 0.5.
E. Representative density is shown for the Lys856 region where B.1.1.529 mutated. The contour level is 5σ.
F. Location of the B.1.1.529 amino acid substitutions. The residues are shown as spheres of their respective Cα-atoms.

**Figure S2.**
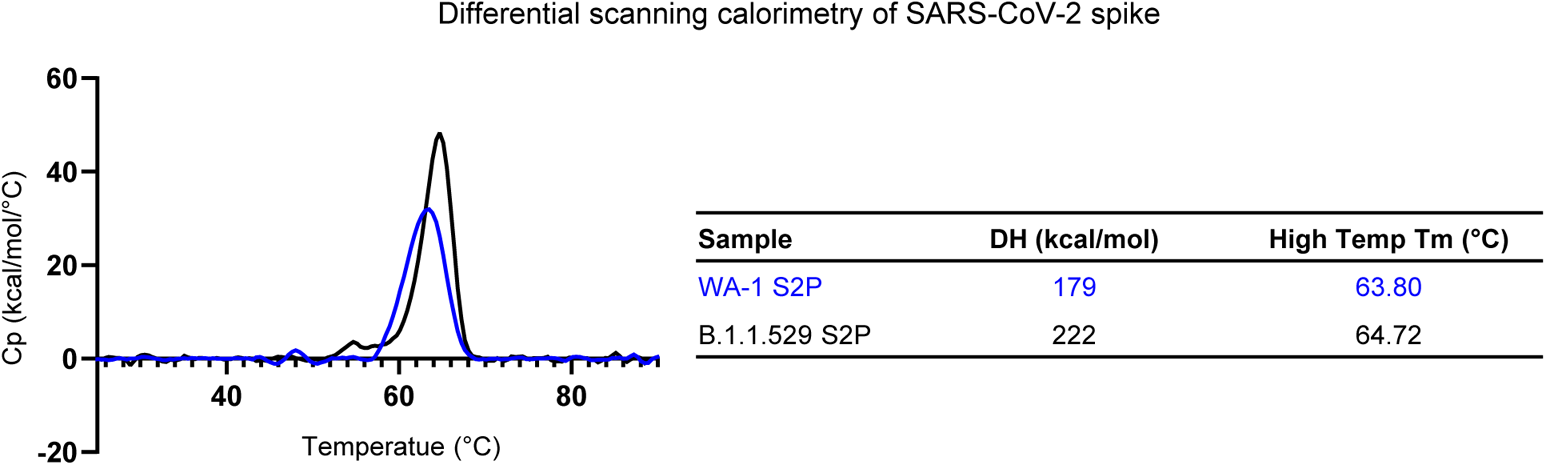
Differential scanning calorimetry of SARS-CoV-2 spikes (S2P) for WA-1 and B.1.1.529 variants.

**Figure S3.**
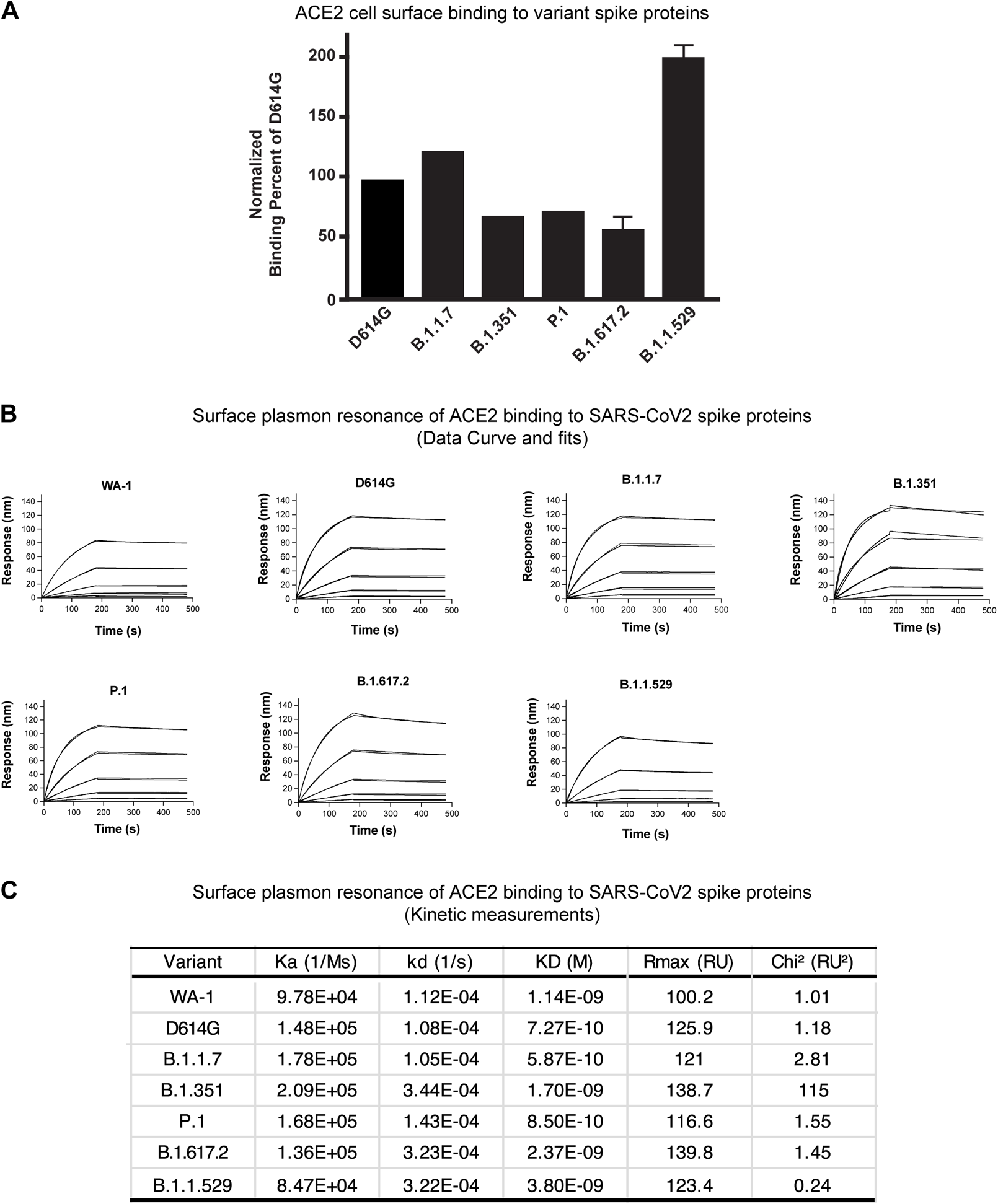
Monomeric ACE2 binding to SARS-CoV-2 B.1.1.529 cell surface expressed spike and S-2P trimer. A. Full length spike proteins from the indicated SARS-CoV-2 variants were expressed on the surface of transiently transfected 293T cells and binding to dimeric human ACE2 was assessed by flow cytometry. Shown is the mean fluorescence intensity (MFI) of bound ACE2 on the indicated cell relative to the MFI of ACE2 bound to D614G expressing cells. The data is expressed as a percentage and shown as the average of three independent experiments.
B. Raw and fitted binding curves for monomeric human ACE2 protein to stabilized Spike 2-proline (S-2P) proteins for each of the indicated variants.
C. Shown are the kinetic parameters, Rmax and Chi^2^ values obtained using the curves in panel B.

**Figure S4.**
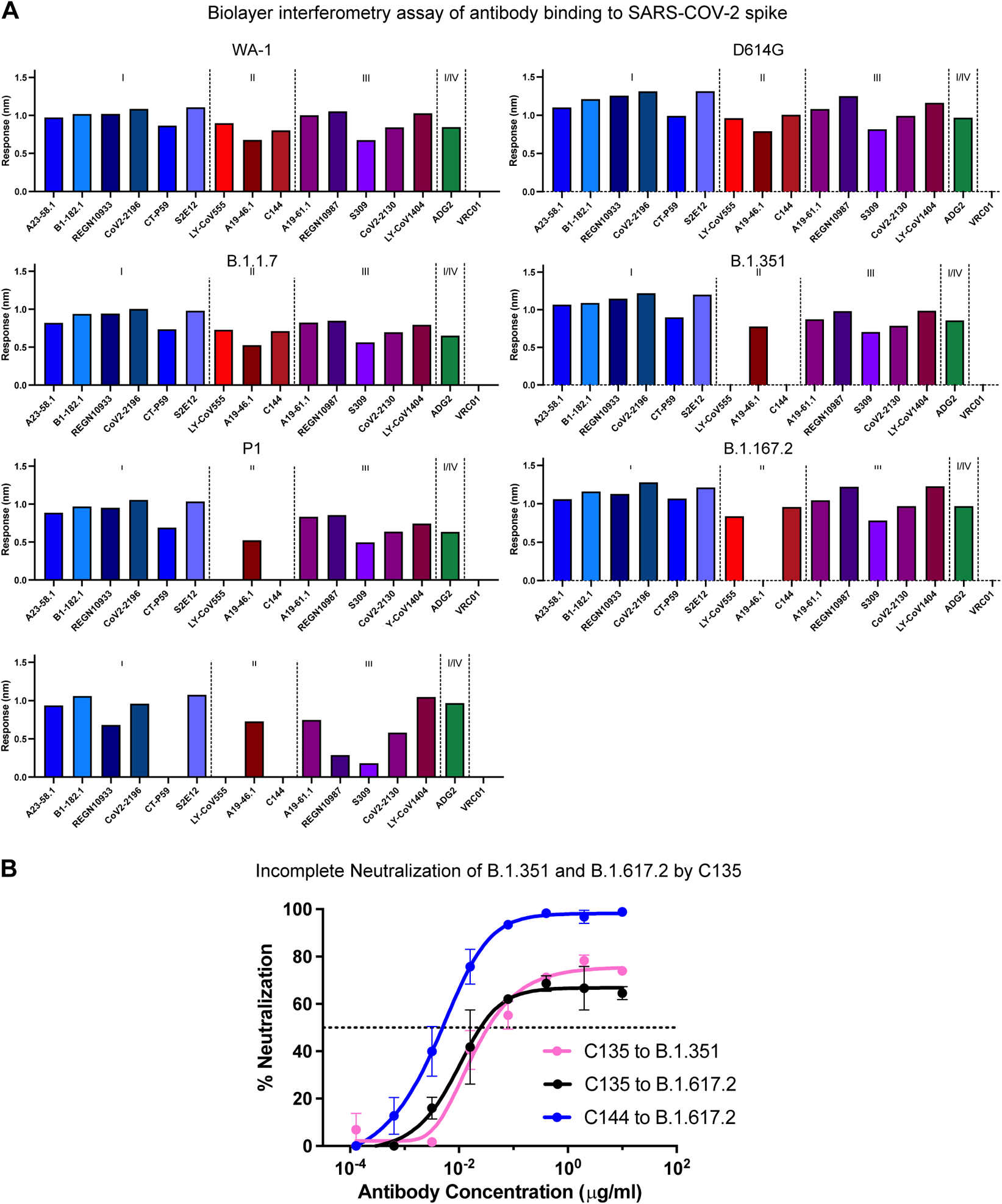
Antibody binding to S2P spike proteins and Neutralization of B.1.351 and B.1.617.2 by C135. A. Biolayer interferometry assay of antibody binding to purified spike proteins of SARS-CoV-2 variants. Antibodies are. Grouped by the Barnes Class I-IV.
B. Lentiviruses pseudotyped with SARS-CoV-2 spike proteins from B.1.351 or B.1.617.2 were incubated with serial dilutions of the indicated concentrations of either C135 or C144 and %neutralization measured.

**Figure S5.**
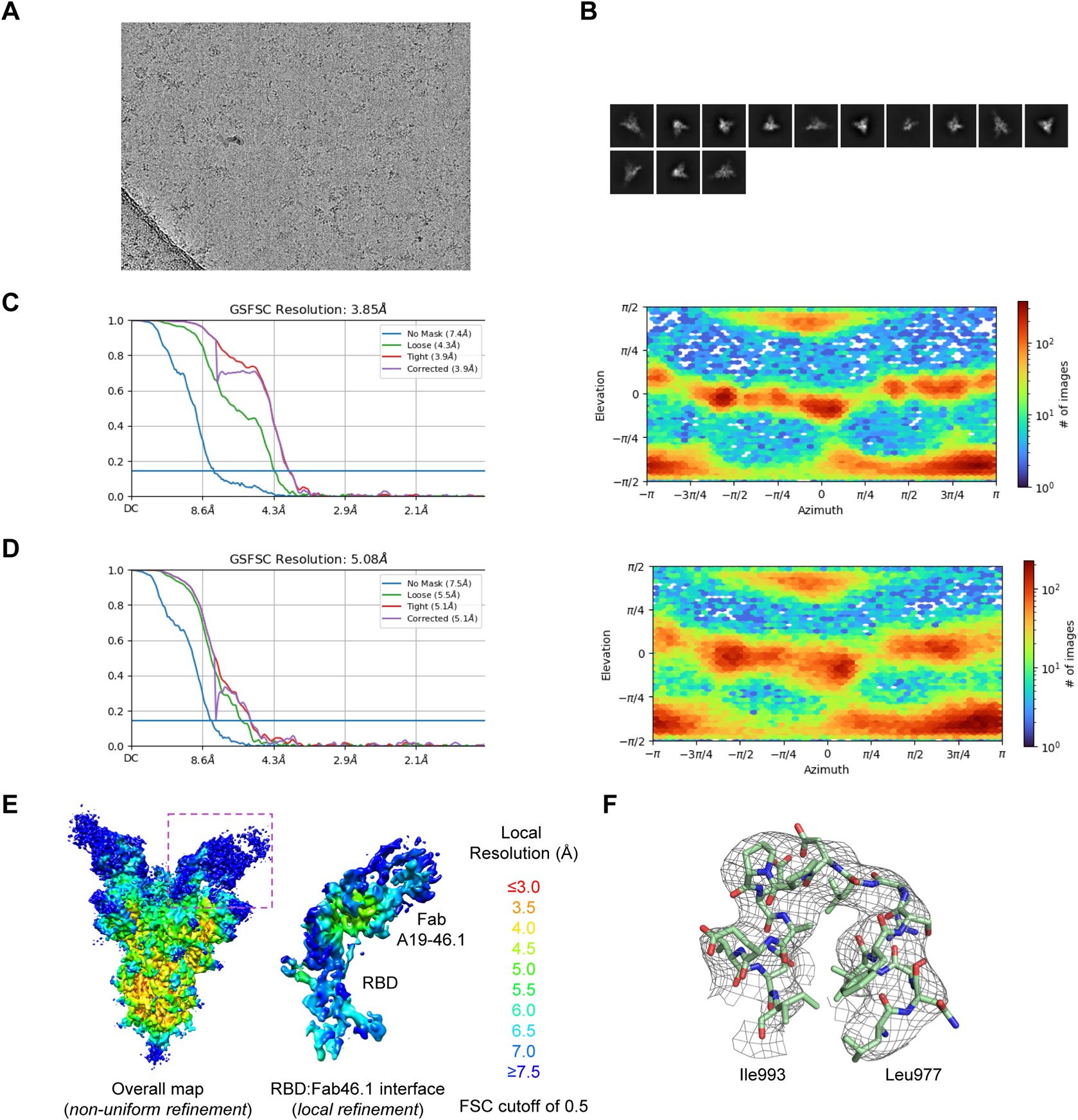
Cryo-EM details of A19-46.1 Fab in complex with SARS-CoV-2 B.1.1.529 spike. A. Representative micrograph.
B. Representative 2D class averages.
C. The gold-standard Fourier shell correlation resulted in a resolution of 3.85 Å for the overall map using non-uniform refinement with C1 symmetry (left panel); the orientations of all particles used in the final refinement are shown as a heatmap (right panel).
D. The gold-standard Fourier shell correlation resulted in a resolution of 5.08 Å for the masked local refinement of the RBD:A19-46.1 interface (left panel) obtained using particle subtraction followed by local refinement; the orientations of all particles used in the local refinement are shown as a heatmap (right panel).
E. The local resolution of the final overall map and locally refined map is shown contoured at 0.068 (4.5σ) and 0.311 (13.9σ), respectively. Resolution estimation was generated through cryoSPARC using an FSC cutoff of 0.5.
F. Representative density is shown for the spike residues 977-993 region. The contour level is 1.5σ.

**Figure S6.**
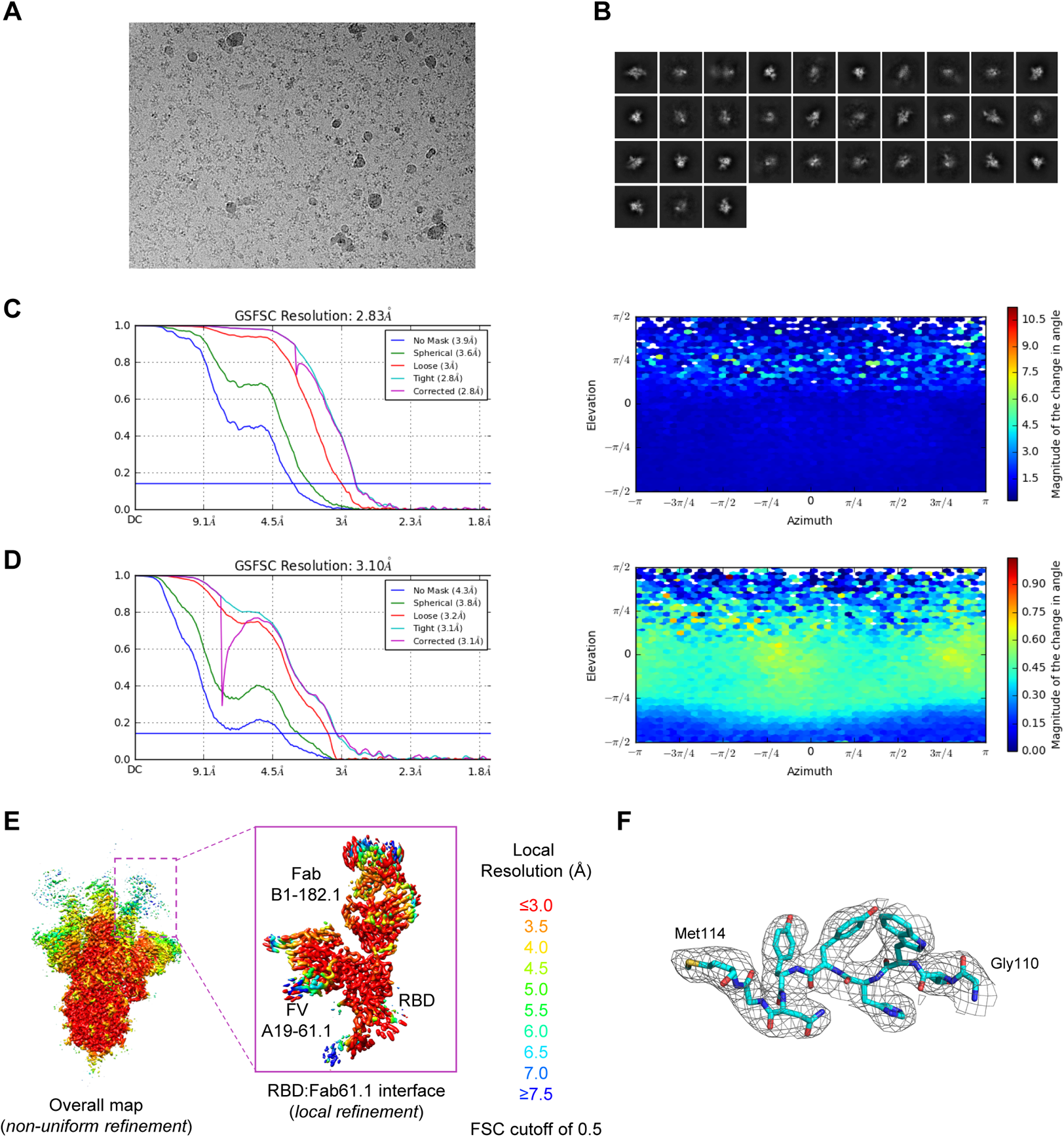
Cryo-EM details of A19-61.1 Fab and Fab B1-182.1 in complex with SARS-CoV-2 WA-1 spike. A. Representative micrograph.
B. Representative 2D class averages are shown.
C. The gold-standard Fourier shell correlation resulted in a resolution of 2.83 Å for the overall map using non-uniform refinement with C1 symmetry (left panel); the orientations of all particles used in the final refinement are shown as a heatmap (right panel).
D. The gold-standard Fourier shell correlation resulted in a resolution of 3.1 Å for the masked local refinement of the RBD:A19-61.1-B1-182.1 ternary interface (left panel) obtained using particle subtraction followed by local refinement; the orientations of all particles used in the local refinement are shown as a heatmap (right panel).
E. The local resolution of the final overall map and locally refined map is shown contoured at 0.154 (5.7σ) and 0.246 (22.2σ), respectively. Resolution estimation was generated through cryoSPARC using an FSC cutoff of 0.5.
F. Representative density is shown for portion of the CDR H3 of A19-61.1 after local refinement. It is evident that the side chains are well defined. The contour level is 10σ.

**Figure S7.**
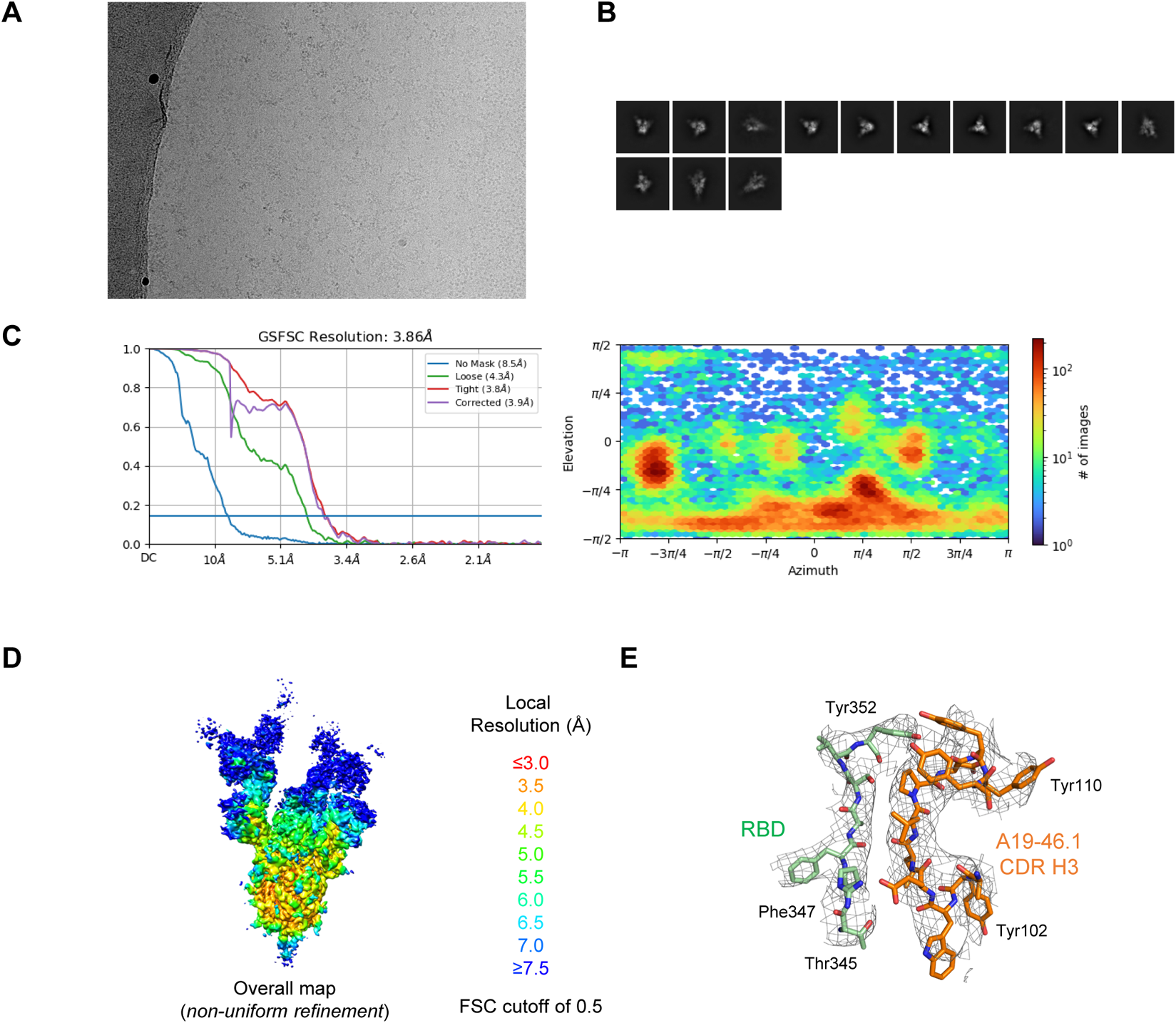
Cryo-EM details of A19-46.1 Fab and Fab B1-182.1 in complex with SARS-CoV-2 B.1.1.529 spike. A. Representative micrograph.
B. Representative 2D class averages are shown.
C. The gold-standard Fourier shell correlation resulted in a resolution of 3.85 Å for the overall map using non-uniform refinement with C1 symmetry (left panel); the orientations of all particles used in the final refinement are shown as a heatmap (right panel).
D. The local resolution of the final overall map is shown contoured at 0.25 (8.5σ). Resolution estimation was generated through cryoSPARC using an FSC cutoff of 0.5.
E. Representative density is shown for B.1.1.529 RBD and Fab 19-46.1 CDR H3 interface. The contour level is 1.5σ.

**Table S1.**
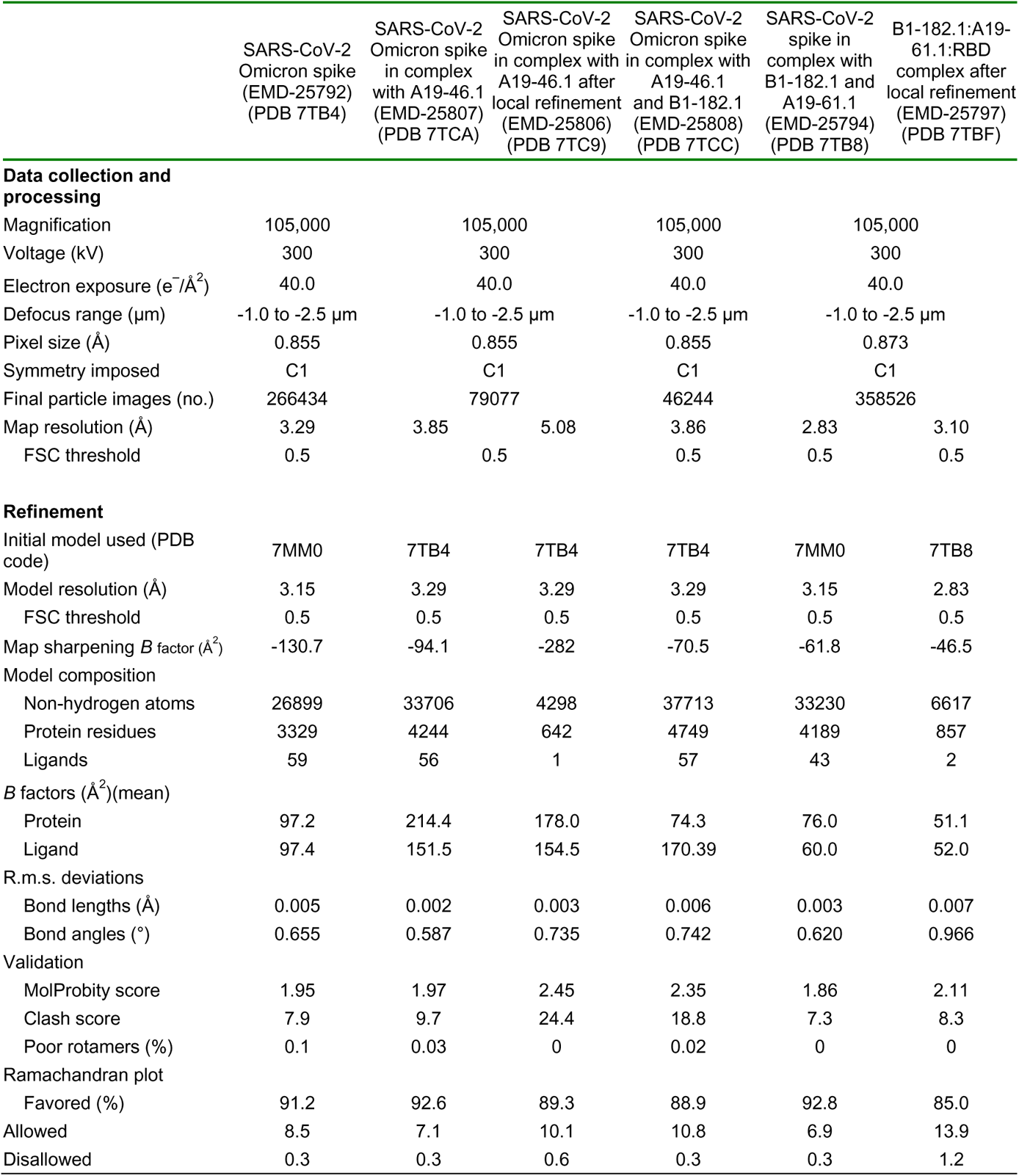
Cryo-EM Data Collection, Refinement and Validation Statistics for SARS COV-2 Spike and Antibody Complexes.

**Table S2.** B.1.1.529 mutation introduced hydrogen bonds and salt bridges.

| Hydrogen bonds |  |  |  |
| --- | --- | --- | --- |
| ## | Protomer 1 | Dist. [Å] | Protomer 2 |
| 1 | C:SER 982[OG | 2.51 | B:LYS 547[NZ] |
| 2 | C:LYS 856[NZ] | 2.60 | B:THR 572[OG1] |
| 3 | B:LYS 764[NZ] | 2.78 | A:GLN 314[OE1] |
| 4 | B:LYS 856[NZ] | 2.99 | A:THR 572[OG1] |
| 6 | C:LYS 547[NZ] | 3.89 | A:ASN 978[OD1] |
| Salt bridges |  |  |  |
| ## | Protomer 1 | Dist. [Å] | Protomer 2 |
| 1 | B:LYS 856[NZ] | 3.94 | A:ASP 568[OD2] |
| 2 | C:ASP 571[OD2] | 3.42 | A:LYS 856[NZ] |

